# Epidemiological model selection unravels the molecular processes of antiviral resistance from experimental phytoplankton-virus dynamics

**DOI:** 10.64898/2026.10.06.757099

**Authors:** Rousseau Raphaël, Bugnot Claire, James Amandin, Yau Sheree, Piganeau Gwenaël, Gourbière Sébastien

## Abstract

Phytoplankton-virus interactions are pivotal in understanding the intertwined dynamics of these key players of the global carbon cycle. Although antiviral resistance is the main hypothesis to solve the apparent paradox of their coexistence, identifying its molecular determinisms and mechanisms, as well as their dynamical consequences, remains challenging. By combining hypotheses about such determinisms and mechanisms, we designed 36 dynamical models and fitted them to experimental infection kinetics in a pioneering selection model approach. Through such a multiple hypotheses testing, we identified that resistance in the model species *Ostreococcus tauri* is most likely to be predominantly intracellular, to control lysis through budding, and to be acquired by phenotypic plasticity. We further showed that viral kinetics are strongly impacted by both the antiviral resistance’s molecular determinants and virions’ inactivation through attachment to lysis detritus, as the latter generates a key negative feedback loop during the ‘bust’ stage of the interaction dynamics.

## 1. Introduction

Phytoplankton are responsible for almost half of the world’s primary production (*1*–*3*) and for the sequestration of 20 to 35% of annual anthropogenic CO_2_ emissions by the global ocean (*4, 5*). Over the past three decades, experimental and field studies of their interactions with lytic viruses have shown the latter to be key engineers of aquatic ecosystems (*6*–*8*). Infections and lysis indeed shape phytoplankton host communities through virus-mediated coexistence (*6, 9*), as stated by the ‘Kill(ing)-the-Winner’ (*10*) and the ‘Boom-and-Busted’ (*11*) hypotheses. Such dynamical interactions influence biogeochemical cycles by facilitating the use of nutrients and carbon by microorganisms within the surface water through the so-called viral shunt (*12*), and their export and long-term storage in the deep waters through the viral shuttle (*13*). To improve our understanding of how molecular processes underlying phytoplankton-virus interactions can shape the ecological impacts of lytic viruses is therefore of paramount importance in the context of accelerating global changes.

Viruses are the most abundant entities in aquatic environments with an average density of ~10^7^ virus-like particles per ml^−1^ (*6*) resulting in an estimated over ~10^23^ infections occurring every second in the global ocean (*8, 14*), which primarily impact unicellular organisms including phytoplankton (*15*). The high efficiency of lytic marine viruses and the lack of temporal or spatial phytoplanktonic host refuges strikingly contrasts with their ability to coexist in both natural environments and experimental settings. A widespread explanation to this apparent paradox relies on the existence of resistant cells that have been characterized in several cyanobacterial (*16, 17*) and eukaryotic (*18*–*26*) host species. Since antiviral resistance is indeed expected to facilitate host-virus coexistence (e.g. (*18, 26*)), there is an obvious need to strengthen our knowledge of the molecular processes of phytoplankton resistance and to identify how they scale up on the population dynamic of phytoplankton-virus interactions.

The production of resistant cells in phytoplankton can be associated with typical determinisms of life-history trait variations; genomic mutations and (induced or random) phenotypic plasticity as previously demonstrated or suggested for other unicellular organisms such as bacteria (*27, 28*), archaea (*29, 30*) and yeast (*31*) Phytoplankton antiviral resistance has been linked to genetic mutations in *Micromonas* (*20, 32, 33*), chromosomal rearrangements in *Ostreococcus* (*26, 34*) and genomic islands in *Prochlorococcus* (*16*). Meanwhile, phenotypic switches in susceptibility/resistance following experimental infections have been shown to correlate with transcriptomic and/or metabolomic changes in *Gephyrocapsa huxleyi* (*35, 36*) and on a specific outlier ‘immunity’ chromosome in *O. tauri* (*37, 38*) and *O. mediterraneus* (*39*).

Besides such determinisms, antiviral resistance can occur through different molecular mechanisms that aim at disrupting different stages of virus transmission and replication (*40*). Extracellular mechanisms limiting viral adsorption or entry into the host cell are thought to be efficient against lytic viruses infecting *M. pusilla* (*20, 33*), *O. tauri* (*37, 38*), *O. mediterraneus* (*39*), *Prochlorococcus* and *Synechococcus* (*16, 17*) and *G. huxleyi* (*21, 36*). When such an extracellular resistance is absent or incomplete, intracellular mechanisms can limit viral replication or control the release of virions into the environment. While the reduction of replication has been proposed to alter the production of virions in *Ostreococcus* (*37, 39*), *Prochlorococcus, Synechococcus* (*17*) and *G. huxleyi* (*36*), the release of viral particles in the environment by budding (rather than by lysing) has also been suggested in so-called ‘resistant producer’ cells (*22, 41, 42*). All the above mechanisms typically aim at preventing infection or lysis, and therefore promote phytoplankton individual survival, despite potential fitness costs (*43*). Meanwhile, it has been suggested that unicellular organisms can also commit ‘altruistic suicide’ by entering a programmed cell death upon infection (*41, 44, 45*). This provides a collective population level defence mechanism by reducing the production of virions and the corresponding force of environmental infection that can complement the above forms of individual level resistance. Given the diversity of individual and collective host defence mechanisms potentially at works, distinguishing their respective contributions in shaping phytoplankton-virus dynamics remains a key challenge, even in the above emblematic host-virus systems where they are under increasing -omic studies scrutiny (*17, 26, 36, 37, 39, 46*).

In this contribution, we intend to develop a comprehensive theoretical framework to quantify the impact of molecular processes of antiviral resistance on the dynamical interactions between phytoplankton and virus. We take advantage of the model system *O. tauri*, an ubiquitous and key ecological species of eukaryotic phytoplankton (class *Mamiellophyceae)* (*47, 48*), and its highly specific double-stranded DNA prasinovirus OtV5 (*49*). We adopt an integrative approach allowing to exploit host and viruses time-series typically provided by experimental evolution of resistance (*18, 35, 41, 50*) through multiple hypotheses testing based on model selection (*51*–*53*). To do so, we aimed at fitting epidemiological models to the dynamical outcomes of *O. tauri*–OtV5 experimental infections. We designed a set of 36 models to represent every combination of the candidate molecular determinisms (genomic mutations, virus-induced or random phenotypic) and mechanisms (extracellular, complete or partial with budding intracellular resistance) of antiviral resistance in the presence or absence of collective defence through programmed cell death. We thereby illustrate how combining experimental infection dynamics and specifically tailored epidemiological models can help identifying the molecular processes underlying phytoplankton-virus interactions.

## 2. Materials and methods

### General overview

We set-up an approach testing multiple hypotheses through model selection that relies on five steps. First, an infection experiment of *O. tauri* by OtV5 provided kinetics data of both hosts and viruses whose abundances were monitored for a period of 20 days. Second, all combinations of hypotheses about the determinisms and mechanisms of antiviral resistance were represented by specifically designed epidemiological models of phytoplankton-virus interactions. Third, for each of the 36 designed models, the predicted dynamics were fitted to the observed variations in *O. tauri* and OtV5 abundances, which provided a Maximum Likelihood Estimate (MLE) for each model parameter. Fourth, the quality of the fits obtained with each model were compared, using Akaike Information Criterion (AIC) (*54*), to identify which of those models and the corresponding hypotheses about the molecular processes of antiviral resistance were best supported by the observed kinetics. Fifth, the predictive capacities of the best models were then assessed using an independent replicate of the *O. tauri* infection by OtV5 produced in the same experimental conditions.

### Infection experiment

#### Cultures and growth conditions

The infection experiment was run with *O. tauri* strain RCC4221 and prasinovirus OtV5 using a L1 medium (NCMA, Bigelow Laboratory for Ocean Sciences, USA) prepared with artificial seawater (ASW) (0.42 M NaCl (S3014, Sigma-Aldrich), 10 mM KCl (P9333, Sigma-Aldrich), 20 mM MgCl_2_ (M2670, Sigma-Aldrich), 10 mM CaCl_2_ (C5080, Sigma-Aldrich), 24.5 mM MgSO_4_ (M5921, Sigma-Aldrich), 2.5 mM NaHCO_3_ (S6297, Sigma-Aldrich)). Four-litre cultures were maintained in 5-litre volume Schott bottles with vented caps and incubated at 20 °C under a 12:12 h light/dark cycle (100 μE m^−2^.s^−1^). The bottles were shaken manually once a day to homogenise the cultures and their position within the incubator was regularly alternated at random to minimise any border effects associated with lighting heterogeneity. Prior to viral inoculation, the main culture batch was first made axenic by single-cell sorting using a flow cytometer (Cytek Aurora), thereby reducing bacterial contamination to less than 1%. A total of 4 independent cultures were established from this main culture batch. Two uninfected controls were maintained for 20 days as references for the susceptible phenotype, while the other two were infected using a concentrated OtV5 lysate at a virus particle to cell ratio of ~1. Cultures were infected 5 days after the start of the phytoplankton experimental growth when cultures were in log phase.

#### Flow cytometry

The cultures were sampled at 4, 8, 14 and 20 hours for a day post-infection (hpi), and then every 24 hours for 14 days. All samples were used to count phytoplankton cells and viral particles by flow cytometry using a Beckman-Coulter Cytoflex equipped with a 488 nm excitation laser. The samples were fixed for 15 minutes in the dark with 0.25% glutaraldehyde (G6257, Sigma-Aldrich) and 0.1% Pluronic F-68 (P-7061, Sigma-Aldrich) (*55*), flash frozen in liquid nitrogen, then stored at −80°C until analysis within a month of sampling. Phytoplankton cells were identified on the basis of their red chlorophyll autofluorescence (> 620 nm) and side scatter (SSC-H). Viral particles were counted based on the protocol of (*56*) where samples were rapidly thawed at 37ºC, then diluted 20–200 fold in < 0.02 μm filtered TE buffer (10 mM Tris-HCl, 1 mM EDTA, pH 8.0) containing SYBR Green I (0.5× final concentration of the commercial stock) and incubated for 10 minutes at 80°C. The trigger was set on green fluorescence (FITC-H) and virus-like particle populations were discriminated on FITC-H intensity and side scatter (SSC-H). Data analysis was performed using CytExpert 2.2 software (Beckman-Coulter).

### Epidemiological modelling of phytoplankton-virus interactions

The 36 designed models all rely on a common Susceptible-Infected-Virus (SIV) core model representing the spread of a pathogen within its host population, to which representation of the hypotheses about the determinisms and mechanisms of resistance were added.

#### Core SIV model

The core model provides a simple description of a unicellular phytoplanktonic host population made of susceptible (S) and infected (I) individuals. Susceptible individuals asexually divide at an intrinsic rate *b* downregulated by intraspecific competition with intensity *c*. Although infected cells still absorb nutrients, and therefore compete with susceptible ones, such resources are considered to be used for viral replication (*57, 58*) and no longer available for cell division. All hosts suffer from intrinsic mortality at rate *d*, and infected cells are further lysed at rate *a*. Upon lysis, *b*_*v*_ new virions are released into the environment and join the pool of viral particles (V) available to produce new infections through contacts with susceptible hosts at rate *β*. Such free environmental virions decay because of deleterious abiotic factors at rate *d*_*v*_. Accordingly, the core SIV model is defined by the following set of equations (Eq. 1-3):

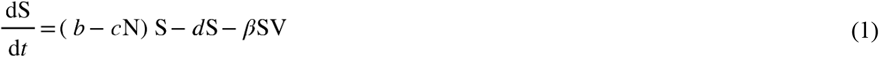

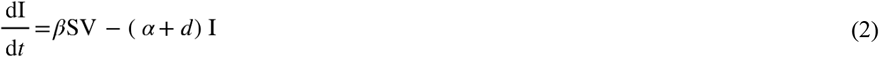

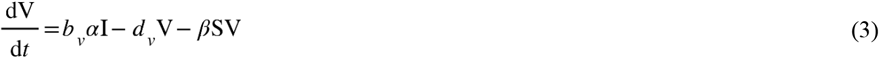

where N = S + I stands for the host population size.

#### Determinisms of antiviral resistance

Antiviral resistant cells can be produced by genomic mutations (GM) or by random (RPP) or virus-induced (IPP) phenotypic plasticity. Under the GM hypothesis, resistant cells emerge according to a mutation probability *μ* during the division of susceptible individuals occurring at an effective rate reduced by the host intraspecific competition (Fig. 1A). By contrast, under each phenotypic plasticity hypothesis, the switch from a susceptible to a resistant phenotype is no longer associated with cell division and a susceptible phenotype can be recovered through reverse gene regulatory changes (Fig. 1B-D). When considering RPP, phenotypic switches can occur throughout the host’s lifetime with respect to acquisition and loss rates γ and γ*’* (Fig. 1B). When resistance is acquired through IPP, acquisition is expected to depend on the extracellular or intracellular mechanism by which such resistance is achieved. In the extracellular case, resistance is triggered at a rate *δ*_*e*_ upon contact between a virus and a susceptible host (Fig. 1C), while in the intracellular case, its induction occurs at the rate *δ*_*i*_ during the early stage of the resulting ‘pseudo-infection’ (Fig. 1D). In both cases, the loss of the resistant phenotype was still assumed to occur at a constant *δ’* (Fig. 1C and D).

**Figure 1.**
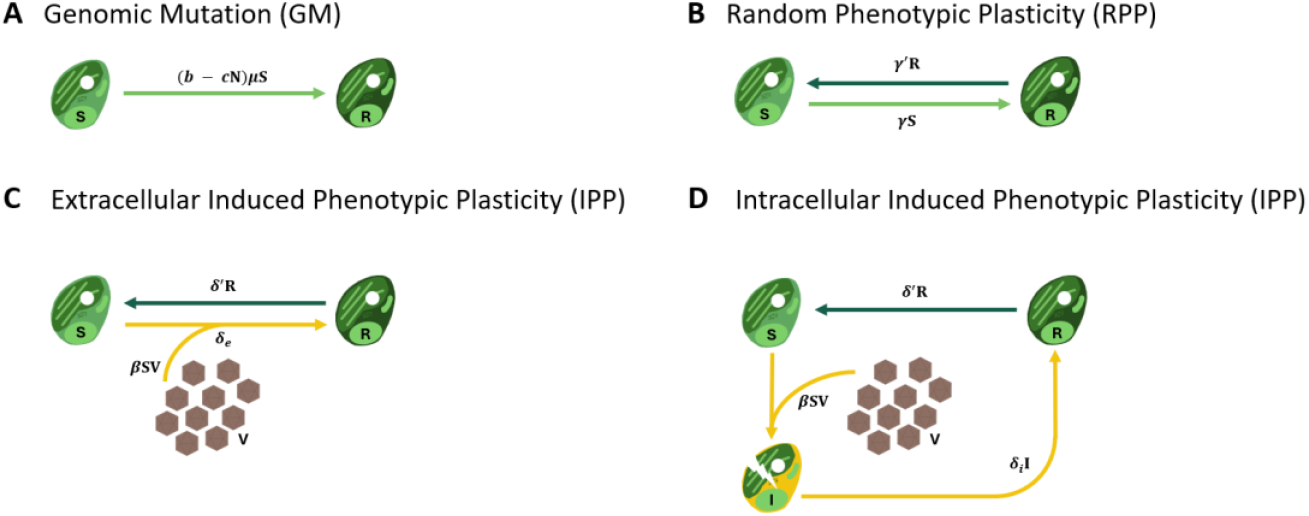
Modelling of the determinism of antiviral resistance in phytoplankton. Resistant cells can be produced through (A) Genomic Mutations, (B) Random Phenotypic Plasticity or (C-D) (Virus-) Induced Phenotypic Plasticity. When resistance is induced by viruses, its rate of acquisition changes according to the extracellular (C) or intracellular (D) mechanism by which resistance is achieved. S, R, I stand for the density of susceptible, resistant, and infected hosts, while V denotes the density of virions in the marine environment. *b, c, β* correspond to the division rate, intensity of intraspecific competition and encounter rate between a susceptible host and a virion, respectively. *μ*, γ, *δ*_*e*_ and *δ*_*i*_ are the probability of acquisition of antiviral resistance by genomic mutation and the rates of random and virally induced phenotypic plasticity. γ*’* and *δ’* are the rates of loss of antiviral resistance through reversed phenotypic changes. Colours are used to discriminate compartment and associated processes; susceptible (light green), resistant (dark green) and infected (yellow) host.

#### Mechanisms of individual and collective host defence

Antiviral individual resistance is achieved through extracellular and/or intracellular host-virus molecular interactions. Extracellular resistance typically reduces the effective encounter rate β^′^ between resistant cells and virions. When such resistance prevents the adsorption of free viral particles, i.e. *β’* = 0, virions were considered to remain unaffected in the marine environment (Fig. 2A). When virions successfully adsorb and enter a resistant host cell; i.e. if *β’* > 0, they can be controlled by intracellular pathways that affect viral replication, neutralize the virus and/or prevent any further release of virions into the environment (Fig. 2B). Meanwhile, such an intracellular control can be partially efficient with so-called ‘resistant producers’ (R^P^) preventing lysis by the release of small quantities of virions through budding at a rate *b*_*vr*_ (Fig. 2C). Such a mechanism of control of infection can potentially lead to a complete clearance, which was considered to occur at a rate *Ψ* and to enable the cell to recover a (non-producer) fully resistant status (R). Collective defence mechanisms rely on the early death of infected hosts to limit viral production and transmission. Such ‘altruistic suicide’ typically involved Programmed Cell Death (PCD) pathways that can be represented by an additional mortality rate *α’* of infected cells (Fig. 2D).

**Figure 2.**
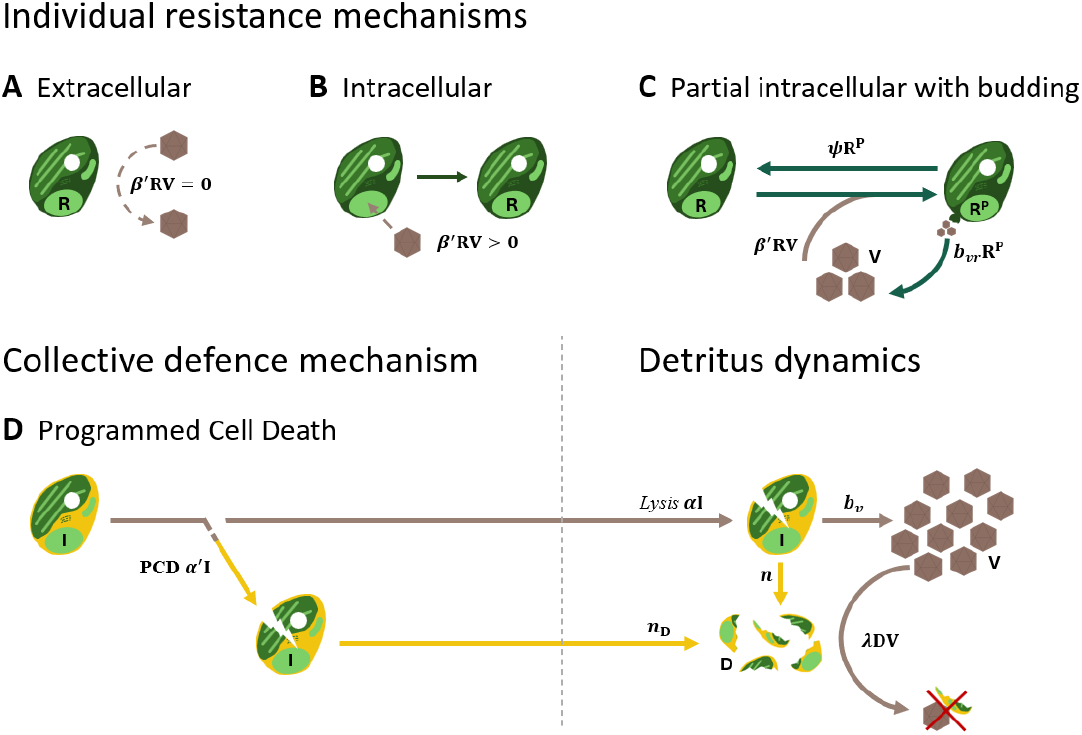
Modelling of the phytoplankton’s individual and collective defence against viruses. Individual resistance can be achieved through (A) extracellular or (B) intracellular molecular mechanisms, or lead to (C) intracellular ‘resistant producers’ preventing lysis through budding, while (D) collective defence involving PCD is provided by the early death of infected individuals. R, R^P^ and I stand for the abundance of non-producer or producer resistant and infected hosts, while V and D denote the abundance of viruses and lysis detritus. *β’* is the encounter rate between a resistant cell and a virus, with *β’* = 0 or *β’* > 0 for extracellular or for (complete and partial with budding) intracellular resistance, respectively. *b*_*v*_ and *b*_*vr*_ stand for the number of virions produced by lysis and budding, and *Ψ* for the rate at which ‘resistant producers’ can clear their partial infection and stop budding. *α* and *α’* are the rate of lysis induced by the virus and the ‘altruistic suicide’ rate of infected individuals. In SRIVD models, each lytic event leads to the release of *n*_D_ detritus onto which environmental virions adsorb at a rate *λ*, so that they are removed from the pool of infectious particles. Colours are used to discriminate compartment and associated processes; resistant (dark green) and infected (yellow) host.

Under any of the above hypotheses about the determinism and mechanisms of phytoplankton antiviral resistance, the demographic parameters of the resistant hosts was identical to that of susceptible cells, except that their birth rate, *b*, was reduced by a typical individual cost ζ of resistance (*43*).

#### Epidemiological models of phytoplankton-virus interactions with antiviral resistance

The SRIV models accounting for resistant cells take on two general forms. The simplest one (Eq. 4-7) describes the phytoplankton-virus interactions in the absence of resistant producer cells;

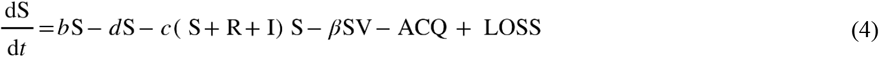

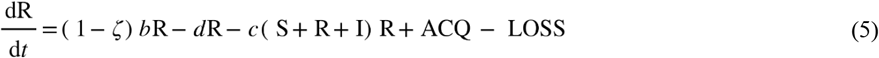

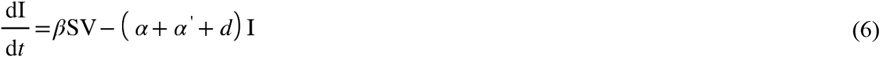

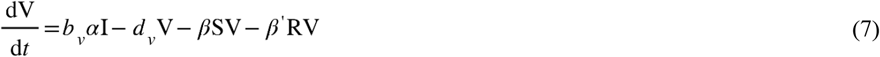

where ACQ and LOSS describe the acquisition and loss of resistance (see SM1) according to each of our 12 combinations of hypotheses about the determinism of resistance (GM, RPP and IPP) and defense mechanisms (extracellular and intracellular, with or without PCD).

The SRIV models accounting for resistant producer cells take on a slightly more complex form (Eq. 8-12) as they include one additional state variable (R^P^);

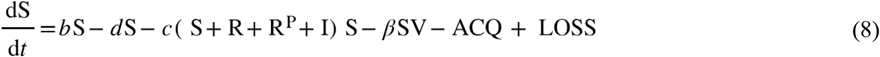

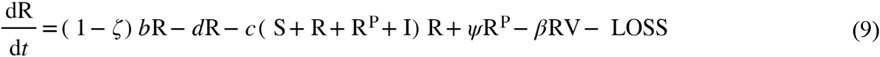

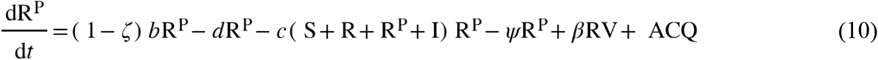

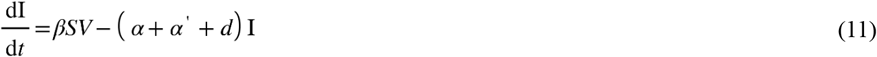

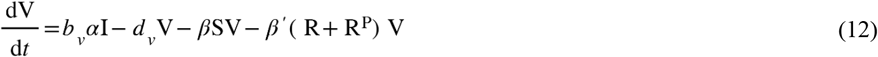

where the definition of the acquisition and loss of resistance terms are presented in SM1 according to each of our 6 additional combinations of hypotheses about the determinism of resistance (GM, RPP and IPP) and this alternative mechanism of partial intracellular resistance associated with budding (with or without PCD).

We further aimed at testing the importance of another identified source of viral mortality on the experimental dynamic of *O. tauri*-OtV5 interaction. Dead cell fragments produced through viral lysis have indeed been suggested to affect the survival and infectivity of virions as they provide decoys for viral adsorption potentially inducing their inactivation through the ejection of their genetic material (*59*–*61*). To predict the significance of such a (dead) host induced viral mortality, we extended each of our SRIV model using a simple description of such interactions with cell fragments. Each lysis event, induced by the virus or an ‘altruistic suicide’, leads to the release of *n*_D_ particles providing non-specific adsorption sites into the environment, which we referred to as detritus. The loss of virions onto such detritus occurs according to a fixed encounter rate *λ* and according to their densities V and D, resulting in the removal of both viruses and detritus from their respective pools as viruses become ineffective and detritus inaccessible for further adsorption.

The 18 SRIV models (with or without R^P^) were therefore expanded into 18 SRIVD models by the addition of i) a viral mortality term (-*λ*DV);

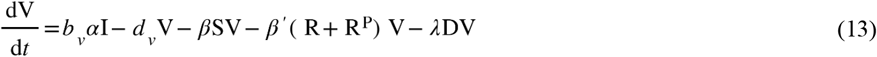

where R^P^ = 0 in the absence of resistant producer cells, and ii) the description of the variations in the amount of lysis detritus D as follows;

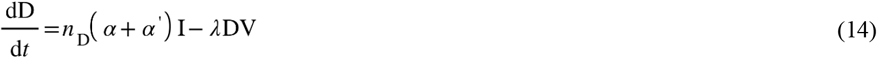

All systems of equations corresponding to those SRIVD models are presented in detail in SM1.

#### Model fitting and parameters estimates

The 36 SRIV and SRIVD models were fitted to the variations in *O. tauri* and OtV5 abundances measured during the first replicate of the infection experiment. Each model *i* (1 ≤ *i* ≤ 36) fitting was achieved by maximizing the following log-likelihood function (Eq. 15);

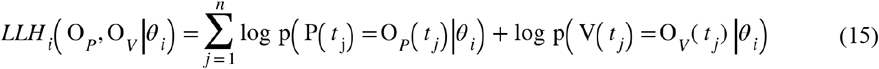

Where P(*t*_*j*_) and V(*t*_*j*_) are statistical variables corresponding to the abundance of phytoplankton and virus expected at time *t*_*j*_ (in hours), O_*P*_(*t*_*j*_) and O_*V*_(*t*_*j*_) stand for the observed abundances at that time of the experiment, and *θ*_*i*_ is the set of parameters corresponding to the model being fitted (see SM1). Probabilities p(P(*t*_*j*_) = O_*P*_(*t*_*j*_)|*θ*_*i*_) and p(V(*t*_*j*_) = O_*V*_(*t*_*j*_)|*θ*_*i*_) were calculated according to negative binomial distributions to account for the overdispersion in phytoplankton and virus densities (*62*) with aggregation parameters *rP* and *rV* estimated separately from *O. tauri* and OtV5 data as described in SM2. The biologically relevant ranges of each parameter of the set *θ*_*i*_ were systematically explored in order to identify the maximum (*L*_*i*_) of the log-likelihood function for a given model *i* and the MLE for each of them.

#### Comparison between models

For each of the 36 models, the maximum likelihood associated with the best fit measures the model inherent ability to describe the experimental data, which rise with the number of parameters (*63*). Models’ comparison was then performed by using the Akaike Information Criterion that allow quantifying when additional parameters actually provide a better representation of the data. Since the complexity-to-data ratio of our most complex model is lower than 40, we used its second-order bias correction, termed AIC_*c*_ (Eq. 16, (*54*)).

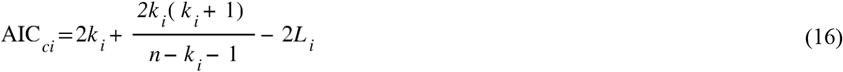

where *K*_*i*_ is the number of parameters of the *i-*th model and *n* is the number of observed data. Accordingly, a better fit, i.e. high (negative) *L*_*i*_, and a lower complexity, i.e. low *k*_*i*_, produce a lower AIC_*c*_. Models were compared using the AIC_*c*_ differences (Δ_*i*_) defined between the AIC_*ci*_ of any model *i* and the lowest AIC_*c*_ obtained for the best model. All models were then ranked according to their ability to fit the data with parsimony, which is typically interpreted as a level of empirical support to the underlying hypotheses (*64*). Such a support is considered to be equivalent to the support received by the best model whenever Δ_*i*_ ≤ 2, considerably lower when 4 ≤ Δ_*i*_ ≤ 7, and essentially null when Δ_*i*_ > 10 (*54*). The AIC_*c*_ differences were also used to calculate the so-called weights of Akaike (*w*_*i*_, Eq. 17)

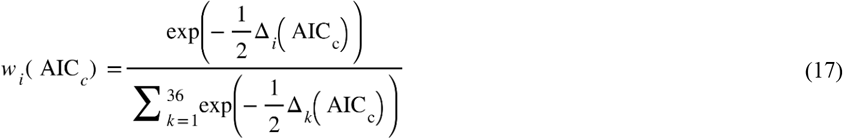

that give the probabilities for each model *i* and the corresponding hypotheses, to be the best representation of the truth (*54*);

#### Assessing the quality and validation of the models

We further assessed the quality of the models fitting and their predictive ability by looking at the post-fitting residual variations and by validation on an independent experimental dataset. We performed linear regression between the predicted and observed values of *O. tauri* and OtV5 abundance using the fitted data and an independent replicate obtained in the same experimental conditions, but from a separate batch of culture. We then used the *R*^2^ to quantify the overall amount of observed variability in the hosts and viruses explained and predicted by the models.

## 3. Results

### Experimental dynamics of *O. tauri*–OtV5 interaction

The *O. tauri* population was grown for 5 days until it reached ~10^7^ cells per ml and before viral inoculation (Figure 3.*a*-*b*). Within a day following the introduction of OtV5 (initial viral particle to cell ratio of ~1), the viral population peaked with a 10 to 12-fold increase in abundance leading to the lysis of > 99% of the host population. In turn, within ~3 days post-inoculation, this drop in host availability led to a sharp decrease in the number of virions, although their abundance remains > 5 times higher than in the inoculum. Soon after this major lytic event, the *O. tauri* population started to recover and rise exponentially, which, as the density of OtV5 remained high, was associated with the production of resistant cells (*65*). The viral density then followed a counter intuitive exponential growth. The concomitant increases in *O. tauri* and OtV5 populations ultimately slowed down to reach a plateau ~11–12 days after inoculation, where their densities stood at ~3.1×10^7^ cells and 1.9×10^8^ virions per ml (i.e. viral particle to cell ratio of ~6.1). The two replicates of this infection experiment provided very similar dynamics (Fig. 3A and B).

**Figure 3.**
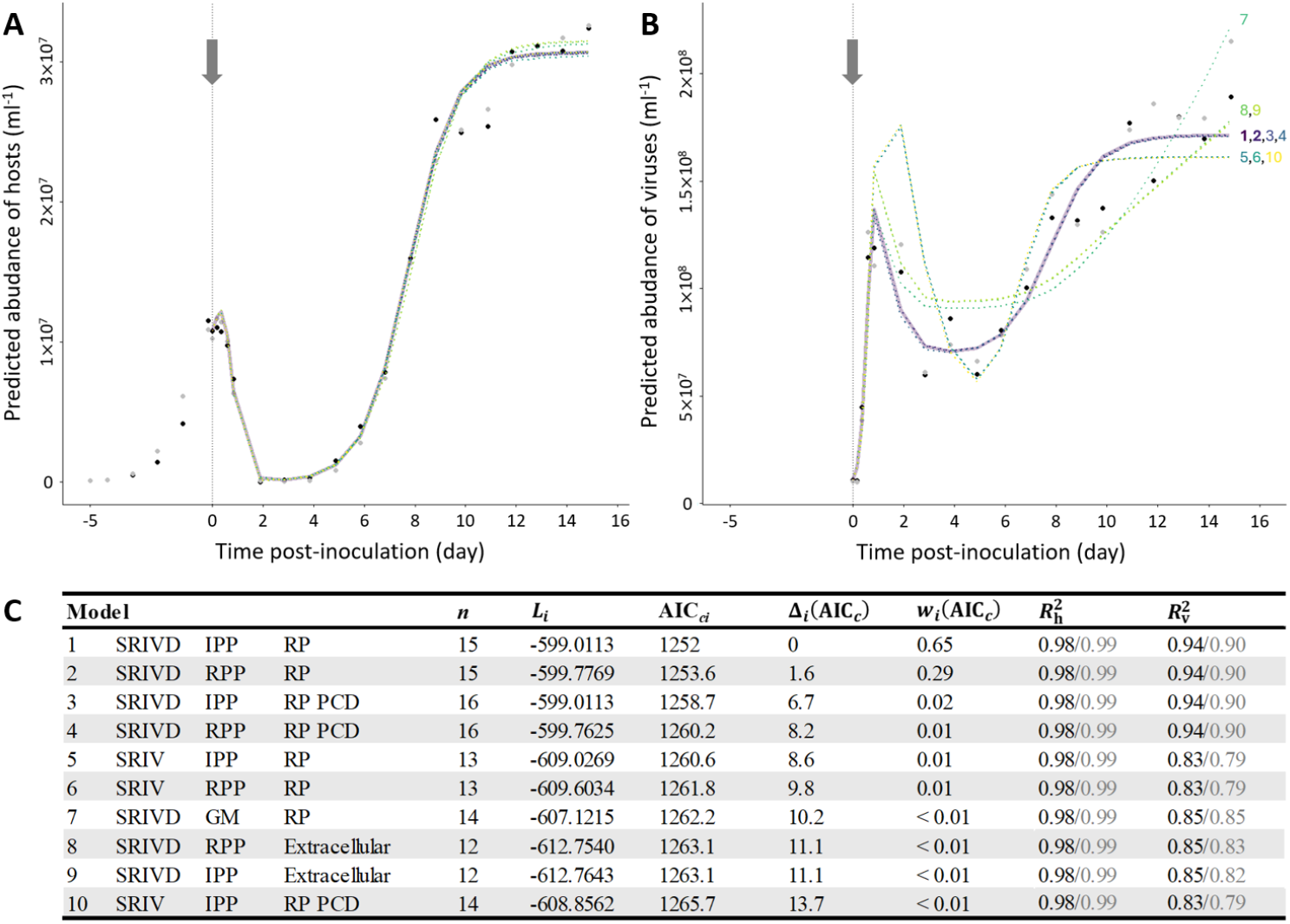
Experimental dynamic of *O. tauri*–OtV5 interaction and phytoplankton-virus model fitting, comparison and validation. (A and B) The variations in *O. tauri* (A) and OtV5 (B) abundances observed in 2 independent replicates that were used to select and validate the models are shown as black and grey points, respectively. The best fits predicted by the top 10 models appear in different colours and numbered according to their ranking (in C). The predictions of the 2 similarly best models and the 8 remaining ones are shown as thick solid and dotted lines, respectively. The grey arrows indicate the point of inoculation. (C) The outcomes of the phytoplankton-virus model fitting, comparison and validation are summarized for the top 10 models ranked according to their merit. Columns provide, for each model, a summary of the hypotheses about the determinism and mechanism of resistance, the number of parameters (*n*), the maximum likelihood (*L*_*i*_) the corrected AIC (AIC_*ci*_), the difference with the AIC_*c*_ of the best model (Δ_*i*_), the weight of Akaike (*w*_*i*_), the coefficients of determination obtained by comparing the observed and predicted abundances of the *O. tauri* hosts and OtV5 virus. In this summary, RP stands for (partial intracellular) Resistant Producers (through budding). Values of *R*^2^ in black and grey correspond to the outcome of the model fitting and validation, respectively.

### Models fitting, comparison and validation

The ability to fit and predict the experimental dynamics were shown to vary substantially between the 36 models (see SM3 for a complete assessment), as evidenced by the results obtained for the top 10 models. Notably, while all those 10 models fit very well the dynamics of *O. tauri* abundance (Fig. 3A), the quality of their adjustment to the OtV5 dynamics is more variable (Fig. 3B). All 10 models indeed explain 98% of the variability observed in phytoplankton abundance (, Fig. 3C) whereas they account for a smaller proportion of the viral dynamics ranging from 83% to 94% (, Fig. 3C). This suggests that signatures of antiviral resistance, and our ability to discriminate between their potential determinisms and mechanisms, are primarily contained in the viral dynamics. The first 4 models, whose weights of Akaike (*w*_i_) indicate that they have a cumulated probability ~0.97 to provide the best representations of the observed dynamics (see Fig. 3C), indeed allow for almost identical fits to the viral dynamics (see Fig. 3B and *L*_*i*_, and in Fig. 3C) as compared to the fits provided by the other top 10 models. Meanwhile, the first and second models were found to be better than the third and fourth, as they provide such a good fit with a smaller number of parameters. The 2 best models then received an equivalent support from the data (as Δ_2_(AIC_*c*_) < 2) while the third and the fourth indeed received a weaker support (Δ_3_(AIC_*c*_) > 4 and Δ_4_(AIC_*c*_) > 7), so that their probability to best represent the observed dynamics (*w*_i_) are 0.65, 0.29, 0.02 and 0.01, respectively.

In addition to their ability to provide very good fits to the dynamic of *O. tauri*–OtV5 interactions, the 4 best models do so for estimates of their parameters that are very consistent with the known host and virus life-histories (Table 1). The MLE of the parameters defining the ‘core SIV’ model (*b, d, c, ζ, β, α, b*_*v*_ and *d*_*v*_) were indeed found both highly similar between the fits of those four best models and close to their previous estimates (Table 1). All 4 models provide an estimate of *O. tauri* population growth rate (i.e. *b* - *d*) equal to 1.1 d^−1^, while the estimated intraspecific competition (*c*) and individual cost of resistance (*ζ*) range from 3.58 to 3.6×10^−8^ d^−1,^ and from 0 to 0.01, respectively. This growth rate lies within its experimentally observed range (0.6 to 1.2) and the corresponding competition intensities produce *O. tauri* densities similar to the ~10^6^–10^7^ cells per ml typically sustained in such batch culture (*22, 38, 66*). Although attempts to measure the cost of resistance for *O. tauri* have yielded variable estimates from different methods, it is commonly found costs to be null or very low (*22, 67, 68*). The estimates of OtV5 lysis rate (*α*) ranging from 3.36 to 3.6 d^−1^ are slightly higher than the previous maximal value found to be around 3 d^−1^, which corresponds to an infection to lysis timing of ~6 to 7 instead of 8 hours. The estimates of the burst size (*b*_*v*_) remain very consistently between 30 and 31, and very close to its previous estimate of 25 (*49*). Similarly, the rate of viral environmental decay (*d*_*v*_) is always lower than 0.013, which consistently matches its estimated value of 0.01 (*69*). Finally, the 4.56–4.8×10^−8^ estimates of the daily rate of infectious encounter per ml (*β*) were found to be about 2 orders of magnitudes below its previous theoretical estimate (*70*), as expected given that the latter was derived from a bio-physical modelling known to quantify the maximal potential of contacts (*71*).

**Table 1.** Experimental and Maximum Likelihood Estimates of the parameters of the 4 best models. ^**(1)**^ From the following formula: *δ*_*i*_ = R(*α*+*α*’+*d*)/(*β*SV-R), assuming the production of 3×10^5^ R in one day following the inoculation at a virus particle to cell ratio of 5 in 3×10^7^ host cells (estimate of *β, α, α*’ and *d* are taken from MLE of Model 1).

| Symbol | Definition | Experimental estimates |  | Maximum Likelihood Estimates |  |  |  | Unit |
| --- | --- | --- | --- | --- | --- | --- | --- | --- |
|  |  | References | Estimate | Model 1 | Model 2 | Model 3 | Model 4 |  |
| $b - d$ | host population growth rate | (67, 72–74) | 0.6 to 1.2 | 1.10 | 1.10 | 1.10 | 1.10 | d<sup>-1</sup> |
| $c$ | intraspecific competition rate | | | $3.60 \times 10^{-8}$ | $3.58 \times 10^{-8}$ | $3.60 \times 10^{-8}$ | $3.58 \times 10^{-8}$ | d<sup>-1</sup> |
| $\zeta$ | individual cost of resistance | (22, 67, 68) | $\sim 0$ | 0 | 0.01 | 0 | 0.01 | 1 |
| $\delta_i$ | rate of resistance acquisition through intracellular IPP | (38)<sup>(1)</sup> | $5.27 \times 10^{-3}$ | $3.60 \times 10^{-3}$ | | $3.60 \times 10^{-3}$ | | d<sup>-1</sup> |
| $\gamma$ | rate of resistance acquisition through RPP | | | | $9.80 \times 10^{-6}$ | | $9.90 \times 10^{-6}$ | d<sup>-1</sup> |
| $\gamma', \delta'$ | rate of resistance loss through phenotypic plasticity | (22, 37) | $< 1.37 \times 10^{-3}$ | $5.20 \times 10^{-10}$ | $4.50 \times 10^{-4}$ | $5.20 \times 10^{-10}$ | $5.90 \times 10^{-4}$ | d<sup>-1</sup> |
| $\Psi$ | virus clearance rate | | | $1.90 \times 10^{-8}$ | 0.016 | $1.90 \times 10^{-8}$ | 0.092 | d<sup>-1</sup> |
| $\beta$ | infectious encounter rate of susceptible hosts | (70) | $1.37 \times 10^{-6}$ | $4.56 \times 10^{-8}$ | $4.56 \times 10^{-8}$ | $4.56 \times 10^{-8}$ | $4.80 \times 10^{-8}$ | ml.d<sup>-1</sup> |
| $\beta'$ | encounter rate of resistant hosts | | | $3.60 \times 10^{-8}$ | $3.60 \times 10^{-8}$ | $3.60 \times 10^{-8}$ | $3.60 \times 10^{-8}$ | ml.d<sup>-1</sup> |
| $\alpha$ | lysis rate | (49, 58, 75) | 1.85 to 3 | 3.60 | 3.60 | 3.60 | 3.36 | d<sup>-1</sup> |
| $\alpha'$ | PCD rate | | | | 0.00 | | 0.029 | d<sup>-1</sup> |
| $b_v$ | burst size | (49) | 25 | 30 | 30 | 30 | 31 | virions.cell<sup>-1</sup> |
| $b_{vr}$ | rate of virions production by budding | (22) | 1 to 3 | 6.24 | 6.24 | 6.24 | 6.24 | d<sup>-1</sup> |
| $d_v$ | decay rate of virions | (69) | 0.01 | 0.013 | 0.012 | 0.013 | 0.00 | d<sup>-1</sup> |
| $\lambda$ | encounter rate of detritus | | | $3.36 \times 10^{-8}$ | $3.36 \times 10^{-8}$ | $3.36 \times 10^{-8}$ | $3.60 \times 10^{-8}$ | ml.d<sup>-1</sup> |
| $n_D$ | number of detritus released by lysis | | | 25 | 25 | 25 | 26 | detritus.cell<sup>-1</sup> |

The ability of the best models to reproduce experimental dynamics for values of the core parameters (*b, d, c, ζ, β, α, b*_*v*_ and *d*_*v*_) consistent with the current knowledge of *O. tauri* and OtV5 life-histories allowed to further compare them to test the typical hypotheses about the determinism and mechanism of antiviral resistance.

### Multiple hypotheses testing

#### Is antiviral resistance determined by genomic mutations, or by a random or induced phenotypic plasticity?

We addressed this question by comparing the models accounting for the emergence of resistant cells through genomic mutations, phenotypic switches induced by viral contact or occurring at a constant rate. The most supported models are by far those considering an *O. tauri* phenotypic plasticity with switches between susceptibility and resistance. All 4 best models indeed make the hypothesis of an IPP or an RPP, and only one of the first 10 models was built on the GM hypothesis. The weights of Akaike associated with the IPP and RPP hypotheses then add up to 0.97 among the 4 best models (and to 0.99 among the first 10 models), which unambiguously identifies phenotypic plasticity as the most likely determinism of *O. tauri* antiviral resistance. The comparison between the IPP and RPP models further showed about twice as much support for a virally induced (*w*_1_ + *w*_3_ = 0.67) than for a random (*w*_2_ + *w*_4_ = 0.30) phenotypic plasticity (and this larger support remains similar among the first 10 models). While the difference between these two alternative causes of phenotypic plasticity appears substantial, it is worth remembering that they still both provide excellent fits to both *O. tauri* and OtV5 dynamics. The fits of both IPP and RPP models provided an opportunity to gain estimates of the specific parameters, i.e. switching rates *δ*_*i*_, γ, *δ*’ and γ’, that remain challenging to assess experimentally. Interestingly, the rate of phenotypic switch toward resistance under the IPP model (*δ*_*i*_, 3.6×10^−3^ d^−1^) is consistent with the estimate that can be derived from a previous *O. tauri* infection experiment (5.27×10^−3^ d^−1^, *(38)* and Table 1), while the rate of such switches under the RPP model is substantially lower (9.8×10^−6^ d^−1^) reinforcing the conclusion that phenotypic switches are more likely to occur through viral induction than at random.

#### Is antiviral resistance extracellular or intracellular, and is it associated with budding?

We answered this convoluted question by comparing models accounting for extracellular or intracellular resistance, and for the release of virions in the environment during chronic infection. Those comparisons clearly show that a partial intracellular resistance associated with budding best reproduces the *O. tauri–*OtV5 interaction dynamics. All 4 best models (and 8 out of the first 10 models) indeed rely on such a resistance and their weights of Akaike add up to 0.97 (and ~0.99 among the top 10 models), which strongly supports the hypothesis of a partial intracellular resistance in *O. tauri*. Interestingly, such a mechanism only provides a partial protection to the phytoplankton population. The estimated effective encounter rate leading to the entry of the virus within such so-called resistant-producer cells (i.e. *β’* = 3.6*×*10^−8^ ml.d^−1^) is indeed obviously larger than under the hypothesis of extracellular resistance (when *β’* = 0), but only ~21% lower than the infectious encounter rate with susceptible cells (*β* = 4.56*×*10^−8^ ml.d^−1^). Meanwhile the rates *Ψ* at a which such cells are able to clear the infection by leaking virions correspond to average recovery time always larger than 11 days, which suggests that *O. tauri* resistant-producer cells are likely to remain infected over their entire lifetime. During this infectious stage, they are estimated to produce about 6 virions d^−1^ per resistant cell (*b*_*vr*_) which is consistent with the unique estimate of this budding production in *O. tauri* (*22*).

#### Do infected cells commit ‘altruistic’ suicide?

The comparison of the models with Programmed Cell Death (hypothesis ‘PCD’) and without such PCD shows almost no support for the existence of ‘altruistic’ suicide during the experimental infection of *O. tauri* by OtV5. Among the first 4 models, the weights of Akaike of the models without PCD adds up to 0.94 (*w*_1_ + *w*_2_), while the weights of those with PCD only represent 0.03 (*w*_3_ + *w*_4_). The third and fourth models (with PCD) indeed provide similar fits as the two best models, which obviously implies that the additional death rate (*α*’) affecting infected cells could not improve the description of the interaction dynamic. The estimates of this parameter tend to be very low, i.e. 0 (model 3) and 0.029 d^−1^ (model 4), reinforcing the conclusion that the protection of *O. tauri* against its highly lytic OtV5 relies on the above mechanisms of individual resistance rather than on a collective defence based on a PCD.

#### Does viral aggregation around lysis detritus shape the dynamics of host-virus interaction?

To detect a potential aggregation of viruses within dead cell fragments produced by host lysis, we compared models with (SRIVD) and without (SRIV) lysis detritus. Interestingly, all 4 top ranked models are SRIVD models (and so are 7 of the first 10 models) so that the sum of their weights of Akaike is therefore equal to 0.97 (and 0.98 among the first 10 models), which strongly support the hypothesis of such OtV5 aggregation. Furthermore, the estimates of the aggregation related parameters obtained for the first four SRIVD models were found to be very consistent, with a 25–26 detritus release per cell lysis event (*n*_D_) and an aggregation rate (λ) ranging in 3.36–3.60*×*10^−8^ ml.d^−1^, which is of the same order of magnitude as the encounter rates between viruses and susceptible (*β*) or resistant (*β’*) hosts. The role played by such an aggregation appears very clearly in the viral dynamics (Figure 3.*b*) since the 3 SRIV models (5, 6 and 10) that lack such feedback fail to reproduce the rapid decline in OtV5 virions following the first major lytic event associated with the release of a large amount of detritus.

## 4. Discussion

Antiviral resistance has been demonstrated in many phytoplankton species (*16*–*26*) and is thought to be essential to explain their coexistence with lytic marine viruses (*18, 26, 76*). Identifying the molecular determinism and mechanisms allowing for such resistance, as well as their impact on phytoplankton–virus population dynamics, therefore stands as a major challenge to understand the dynamic of the global ocean and the ecosystem services it provides. While omics approaches are starting to uncover the molecular bases of such resistance in a few biological systems (*17, 26, 36, 37, 39, 46*), the use of experimental time-series of phytoplankton-virus interaction dynamics, that are being produced on a large number of biological systems, could provide an efficient alternative for model selection to identify these mechanisms from their population dynamics signatures. Here, we used this innovative approach using *O. tauri–*OtV5 experimental dynamics to select among hypotheses about the molecular determinism and mechanisms of antiviral resistance represented through a set of 36 specifically designed host-virus epidemiological models.

The first main outcome of this study is that antiviral resistance has to be associated with phenotypic plasticity to reproduce the observed dynamics. While genomic mutations could allow for the control of the major lytic event, they did not allow for the following swift stabilization of the *O. tauri*–OtV5 interaction dynamics. Meanwhile, phenotypic switches between susceptibility and resistance could convincingly lead to such a stable coexistence and they were found twice more likely to do so while being induced by viruses than when occurring at random. The strong support (i.e. over 99%) to a phenotypically plastic antiviral resistance is consistent with previous transcriptomic comparisons made between susceptible and resistant cells in *Ostreococcus* species (*37*–*39, 75*). Significant differential gene expressions have indeed been identified between susceptible and resistant *O. tauri* lines, most of them occurring on a small outlier chromosome (SOC) (*37*). Meanwhile, recent studies on the closely related species *O. mediterraneus* demonstrated similar transcription changes (still localised on the SOC) between susceptible and resistant lines (*39*). Interestingly, concomitant comparative genomic analyses suggested that the transcriptional changes observed in *O. tauri* could be associated with structural variations on the SOC (*38, 77*), although no chromosomal rearrangements were linked to resistance in 5 independent comparisons of susceptible/resistant lines in *O. mediterraneus* (*78*). While the link between structural variations and gene regulatory changes still remains to be elucidated across *Ostreococcus* species, the outcome of this study suggests that in *O. tauri* the resulting individual phenotypic switch toward resistance has to occur at a rate equivalent to 3.6×10^−3^ per day, which is consistent with previous experimental data (*38*). Meanwhile, the reversed phenotypic switch, from a resistant to a susceptible cell, were predicted to occur at much lower rates close to 5×10^−10^ per day, which concur with the observation that *O. tauri* cells can remain resistant for several years in laboratory cultures (*22, 37*).

The second key outcome of our study is that the protection of *O. tauri* population from their lytic viruses does not involve a collective defence mechanism achieved through programmed cell death and is instead associated with partial intracellular resistance preventing lysis through budding. While ‘altruistic’ suicide has been discussed for other unicellular organisms, including phytoplanktonic species (*41, 44, 45, 79*), we did not find any evidence that it was involved in the control of the OtV5 spread through *O. tauri* population. Instead, the hypothesis of a partial intracellular resistance was strongly supported by our model comparisons and selection. The production of virions by budding was then able to sustain the viral population after a stable coexistence was reached, in agreement with the previous observation of *O. tauri* ‘resistant producers’ cells with partial intracellular resistance (*22*). Interestingly, the proposed alternative hypothesis considering that the production of virions in resistant lineages could be attributed to the lysis of a small sub-population of cells losing their resistance through phenotypic switches (*37, 68*), was represented by models accounting for either extracellular or intracellular resistance mechanisms, but provided fits 2 to 3 orders of magnitude worse. The partial resistance that best fitted the experimental dynamics not only relied on intracellular control of the replication and release of virions, but also on a 21% reduction in the adsorption of virions onto the cells. Such a partial extracellular resistance was indeed observed in another resistant producer line following an adsorption assay (*22*), as well as in other non-producer lines of the *Ostreococcus* genus (*38, 39*). Taken together, the findings echo the main conclusion of a previous theoretical contribution about models of *O. tauri*–OtV5 interaction considering similar molecular processes of antiviral resistance, which showed that their long-term coexistence is most likely under the hypothesis of an admixture of extracellular and intracellular mechanisms (*70*). We must, however, remain cautious in generalising such conclusions to other phytoplankton species, since different type of antiviral resistances have been found within the same strain of *O. tauri* (*22, 37*) as well as in other species such as *G. huxleyi*–EhV, even if the latter also support that some line produced virions by budding (*41*).

A last outcome of our analyses is to provide the first quantitative evidence of the pivotal role of lysis detritus in controlling the temporal dynamic of the number of virions. The models accounting for the aggregation of virions on such detritus indeed received a strong support with a 97% likelihood of representing the natural processes underlying the observed dynamics. Interestingly, this came as such aggregation allows for a rapid drop of viral particles following the major lytic event, which appear to be a typical feature of the dynamics exhibited during infection experiments achieved on several phytoplankton-virus system (*18, 35, 41, 76, 80, 81*). While the impact of such detritus might have been overlooked, the role of sticky polymers, such as transparent exopolymer or TEP, has received more attention. Viruses and phytoplankton cells can aggregate with TEP (*41, 81*–*84*) and sink, so that they are thought to play a major role in the carbon export toward deep seas through the ‘viral shuttle’ (*85*). Meanwhile, a significant difference between detritus and TEP is that aggregation onto lysis detritus increases viruses mortality, while viruses are thought to survive within sticky aggregate (creating potential hotspot of infection (*83*)). Estimating the importance of those two forms of non-specific viral adsorption would thus undoubtedly improve our understanding of marine virus dynamics. The production of lysis detritus being widespread, their integration into ecosystemic modelling (e.g. (*86, 87*)) might help improving our ability to predict ocean carbon fluxes.

This contribution provides the first model selection approach based on fitting the dynamics of phytoplankton-virus observed in experimental infection dynamics and allowed to demonstrate its potential to inform about the most likely molecular processes underlying antiviral resistance by exploiting their population dynamics signatures. Applying this approach to similar time-series in other phytoplankton–virus species could therefore prove to be an efficient strategy to uncover the diversity of mechanisms involved in the resistance of those marine hosts to their lytic viruses. Since this study clearly revealed that the viral kinetic is instrumental in differentiating between models and hypotheses about the fine molecular processes involved in such interactions, future experimental designs would undoubtedly benefit from more frequent and precise monitoring of virions abundance.

## Acknowledgements

We would like to thank Christophe Salmeron and David Pecqueur for their help with the cytometer on the BIOPIC platform.

## Data and materials availability

All data needed to evaluate the conclusions in the paper are present in the paper and/or the Supplementary Materials.

## References

1. C. B. Field, M. J. Behrenfeld, J. T. Randerson, P. Falkowski, Primary Production of the Biosphere: Integrating Terrestrial and Oceanic Components. Science 281, 237–240 (1998).

2. E. Litchman, P. De Tezanos Pinto, K. F. Edwards, C. A. Klausmeier, C. T. Kremer, M. K. Thomas, Global biogeochemical impacts of phytoplankton: a trait-based perspective. J. Ecol. 103, 1384–1396 (2015).

3. A. Z. Worden, M. J. Follows, S. J. Giovannoni, S. Wilken, A. E. Zimmerman, P. J. Keeling, Rethinking the marine carbon cycle: Factoring in the multifarious lifestyles of microbes. Science 347, 1257594 (2015).

4. R. A. Houghton, Balancing the Global Carbon Budget. Annu. Rev. Earth Planet. Sci. 35, 313–347 (2007).

5. S. Khatiwala, F. Primeau, T. Hall, Reconstruction of the history of anthropogenic CO2 concentrations in the ocean. Nature 462, 346–349 (2009).

6. S. W. Wilhelm, A. R. Matteson, Freshwater and marine virioplankton: a brief overview of commonalities and differences. Freshw. Biol. 53, 1076–1089 (2008).

7. S. Jacquet, T. Miki, R. Noble, P. Peduzzi, S. Wilhelm, Viruses in aquatic ecosystems: important advancements of the last 20 years and prospects for the future in the field of microbial oceanography and limnology. Adv. Oceanogr. Limnol. 1, 97–141 (2010).

8. K. D. A. Mojica, C. P. D. Brussaard, Viruses of Plankton: On the Edge of the Viral Frontier. Microorganisms 12, 31 (2023).

9. C. P. D. Brussaard, Viral Control of Phytoplankton Populations—a Review1. J. Eukaryot. Microbiol. 51, 125–138 (2004).

10. T. F. Thingstad, Elements of a theory for the mechanisms controlling abundance, diversity, and biogeochemical role of lytic bacterial viruses in aquatic systems. Limnol. Oceanogr. 45, 1320–1328 (2000).

11. K. J. Flynn, A. Mitra, W. H. Wilson, S. A. Kimmance, D. R. Clark, A. Pelusi, L. Polimene, ‘Boom-and-busted’ dynamics of phytoplankton–virus interactions explain the paradox of the plankton. New Phytol. 234, 990–1002 (2022).

12. S. W. Wilhelm, C. A. Suttle, Viruses and Nutrient Cycles in the Sea. BioScience 49, 781–788 (1999).

13. M. G. Weinbauer, Ecology of prokaryotic viruses. FEMS Microbiol. Rev. 28, 127–181 (2004).

14. C. A. Suttle, Marine viruses — major players in the global ecosystem. Nat. Rev. Microbiol. 5, 801–812 (2007).

15. K. E. Wommack, R. R. Colwell, Virioplankton: Viruses in Aquatic Ecosystems. Microbiol. Mol. Biol. Rev. 64, 69–114 (2000).

16. S. Avrani, O. Wurtzel, I. Sharon, R. Sorek, D. Lindell, Genomic island variability facilitates Prochlorococcus–virus coexistence. Nature 474, 604–608 (2011).

17. S. Zborowsky, D. Lindell, Resistance in marine cyanobacteria differs against specialist and generalist cyanophages. Proc. Natl. Acad. Sci. 116, 16899–16908 (2019).

18. R. Thyrhaug, A. Larsen, T. Thingstad, G. Bratbak, Stable coexistence in marine algal host-virus systems. Mar. Ecol. Prog. Ser. 254, 27–35 (2003).

19. K. Tarutani, K. Nagasaki, M. Yamaguchi, Virus adsorption process determines virus susceptibility in Heterosigma akashiwo (Raphidophyceae). Aquat. Microb. Ecol. 42, 209–213 (2006).

20. A. Zingone, F. Natale, E. Biffali, M. Borra, G. Forlani, D. Sarno, Diversity in morphology, infectivity, molecular characteristics and induced host resistance between two viruses infecting Micromonas pusilla. Aquat. Microb. Ecol. 45, 1–14 (2006).

21. M. Frada, I. Probert, M. J. Allen, W. H. Wilson, C. De Vargas, The “Cheshire Cat” escape strategy of the coccolithophore Emiliania huxleyi in response to viral infection. Proc. Natl. Acad. Sci. 105, 15944–15949 (2008).

22. R. Thomas, N. Grimsley, M. Escande, L. Subirana, E. Derelle, H. Moreau, Acquisition and maintenance of resistance to viruses in eukaryotic phytoplankton populations. Environ. Microbiol. 13, 1412–1420 (2011).

23. K. Kimura, Y. Tomaru, Coculture with marine bacteria confers resistance to complete viral lysis of diatom cultures. Aquat. Microb. Ecol. 73, 69–80 (2014).

24. T. V. Johannessen, G. Bratbak, A. Larsen, H. Ogata, E. S. Egge, B. Edvardsen, W. Eikrem, R.-A. Sandaa, Characterisation of three novel giant viruses reveals huge diversity among viruses infecting Prymnesiales (Haptophyta). Virology 476, 180–188 (2015).

25. C. F. Quispe, A. Esmael, O. Sonderman, M. McQuinn, I. Agarkova, M. Battah, G. A. Duncan, D. D. Dunigan, T. P. L. Smith, C. De Castro, I. Speciale, F. Ma, J. L. Van Etten, Characterization of a new chlorovirus type with permissive and non-permissive features on phylogenetically related algal strains. Virology 500, 103–113 (2017).

26. S. Yau, M. Krasovec, L. F. Benites, S. Rombauts, M. Groussin, E. Vancaester, J.-M. Aury, E. Derelle, Y. Desdevises, M.-L. Escande, N. Grimsley, J. Guy, H. Moreau, S. Sanchez-Brosseau, Y. Van De Peer, K. Vandepoele, S. Gourbiere, G. Piganeau, Virus-host coexistence in phytoplankton through the genomic lens. Sci. Adv. 6, eaay2587 (2020).

27. J. J. Bull, C. S. Vegge, M. Schmerer, W. N. Chaudhry, B. R. Levin, Phenotypic Resistance and the Dynamics of Bacterial Escape from Phage Control. PLoS ONE 9, e94690 (2014).

28. J. M. Broniewski, S. Meaden, S. Paterson, A. Buckling, E. R. Westra, The effect of phage genetic diversity on bacterial resistance evolution. ISME J. 14, 828–836 (2020).

29. L. A. Marraffini, E. J. Sontheimer, CRISPR interference: RNA-directed adaptive immunity in bacteria and archaea. Nat. Rev. Genet. 11, 181–190 (2010).

30. G. Vestergaard, R. A. Garrett, S. A. Shah, CRISPR adaptive immune systems of Archaea. RNA Biol. 11, 156–167 (2014).

31. S. Chau, J. Gao, A. J. Diao, S. B. Cao, A. Azhieh, A. R. Davidson, M. D. Meneghini, Diverse yeast antiviral systems prevent lethal pathogenesis caused by the L-A mycovirus. Proc. Natl. Acad. Sci. 120, e2208695120 (2023).

32. R. E. Waters, A. T. Chan, Micromonas pusilla Virus: the Virus Growth Cycle and Associated Physiological Events Within the Host Cells; Host Range Mutation. J. Gen. Virol. 63, 199–206 (1982).

33. A. Bedi De Silva, S. W. Polson, C. R. Schvarcz, G. F. Steward, K. F. Edwards, Transient, context-dependent fitness costs accompanying viral resistance in isolates of the marine microalga Micromonas sp. (class Mamiellophyceae). Environ. Microbiol. 26, e16686 (2024).

34. S. Yau, C. Hemon, E. Derelle, H. Moreau, G. Piganeau, N. Grimsley, A Viral Immunity Chromosome in the Marine Picoeukaryote, Ostreococcus tauri. PLOS Pathog. 12, e1005965 (2016).

35. M. J. Frada, S. Rosenwasser, S. Ben-Dor, A. Shemi, H. Sabanay, A. Vardi, Morphological switch to a resistant subpopulation in response to viral infection in the bloom-forming coccolithophore Emiliania huxleyi. PLOS Pathog. 13, e1006775 (2017).

36. T. S. Shaler, A. Fromm, D. Schatz, S. Ben-Dor, E. Feldmesser, A. Vardi, Mapping the transcriptional landscape of algal resistance to viral infection reveals a core expression program. New Phytol. 248, 1368–1384 (2025).

37. S. Yau, C. Hemon, E. Derelle, H. Moreau, G. Piganeau, N. Grimsley, A Viral Immunity Chromosome in the Marine Picoeukaryote, Ostreococcus tauri. PLOS Pathog. 12, e1005965 (2016).

38. S. Yau, G. Caravello, N. Fonvieille, É. Desgranges, H. Moreau, N. Grimsley, Rapidity of Genomic Adaptations to Prasinovirus Infection in a Marine Microalga. Viruses 10, 441 (2018).

39. A. James, R. Marcellin-Gros, S. Yau, D. Stien, G. Piganeau, Metabolomic and Transcriptomic rewiring between virus resistance and susceptibility in Ostreococcus mediterraneus. Microbiology [Preprint] (2026). 10.64898/2026.02.10.705097.

40. R. Thomas, S. Jacquet, N. Grimsley, H. Moreau, Strategies and mechanisms of resistance to viruses in photosynthetic aquatic microorganisms. Adv. Oceanogr. Limnol. 3, 1–15 (2012).

41. F. Vincent, U. Sheyn, Z. Porat, D. Schatz, A. Vardi, Visualizing active viral infection reveals diverse cell fates in synchronized algal bloom demise. Proc. Natl. Acad. Sci. 118, e2021586118 (2021).

42. J. Wang, L. Li, S. Lin, Active viral infection during blooms of a dinoflagellate indicates dinoflagellate-viral co-adaptation. Appl. Environ. Microbiol. 89, e01156–23 (2023).

43. C. Winter, T. Bouvier, M. G. Weinbauer, T. F. Thingstad, Trade-Offs between Competition and Defense Specialists among Unicellular Planktonic Organisms: the “Killing the Winner” Hypothesis Revisited. Microbiol. Mol. Biol. Rev. 74, 42–57 (2010).

44. K. D. Bidle, P. G. Falkowski, Cell death in planktonic, photosynthetic microorganisms. Nat. Rev. Microbiol. 2, 643–655 (2004).

45. K. D. Bidle, L. Haramaty, J. Barcelos E Ramos, P. Falkowski, Viral activation and recruitment of metacaspases in the unicellular coccolithophore, Emiliania huxleyi. Proc. Natl. Acad. Sci. 104, 6049–6054 (2007).

46. G. Schleyer, C. Kuhlisch, C. Ziv, S. Ben-Dor, S. Malitsky, D. Schatz, A. Vardi, Lipid biomarkers for algal resistance to viral infection in the ocean. Proc. Natl. Acad. Sci. 120, e2217121120 (2023).

47. M. Tragin, D. Vaulot, Green microalgae in marine coastal waters: The Ocean Sampling Day (OSD) dataset. Sci. Rep. 8, 14020 (2018).

48. M. B. Cooper, E. Kazamia, K. E. Helliwell, U. J. Kudahl, A. Sayer, G. L. Wheeler, A. G. Smith, Cross-exchange of B-vitamins underpins a mutualistic interaction between Ostreococcus tauri and Dinoroseobacter shibae. ISME J. 13, 334–345 (2019).

49. E. Derelle, C. Ferraz, M.-L. Escande, S. Eychenié, R. Cooke, G. Piganeau, Y. Desdevises, L. Bellec, H. Moreau, N. Grimsley, Life-Cycle and Genome of OtV5, a Large DNA Virus of the Pelagic Marine Unicellular Green Alga Ostreococcus tauri. PLoS ONE 3, e2250 (2008).

50. N. Joffe, C. Kuhlisch, G. Schleyer, N. S. Ahlers, A. Shemi, A. Vardi, Cell-to-cell heterogeneity drives host–virus coexistence in a bloom-forming alga. ISME J. 18, wrae038 (2024).

51. J. B. Johnson, K. S. Omland, Model selection in ecology and evolution. Trends Ecol. Evol. 19, 101–108 (2004).

52. N. T. Hobbs, R. Hilborn, Alternatives To Statistical Hypothesis Testing In Ecology: A Guide To Self Teaching. Ecol. Appl. 16, 5–19 (2006).

53. A. T. Tredennick, G. Hooker, S. P. Ellner, P. B. Adler, A practical guide to selecting models for exploration, inference, and prediction in ecology. Ecology 102, e03336 (2021).

54. K. P. Burnham, D. R. Anderson, Multimodel Inference: Understanding AIC and BIC in Model Selection. Sociol. Methods Res. 33, 261–304 (2004).

55. D. Marie, F. Rigaut-Jalabert, D. Vaulot, An improved protocol for flow cytometry analysis of phytoplankton cultures and natural samples. Cytometry A 85, 962–968 (2014).

56. C. P. D. Brussaard, Optimization of Procedures for Counting Viruses by Flow Cytometry. Appl. Environ. Microbiol. 70, 1506–1513 (2004).

57. D. Maat, C. Brussaard, Both phosphorus- and nitrogen limitation constrain viral proliferation in marine phytoplankton. Aquat. Microb. Ecol. 77, 87–97 (2016).

58. A. Monier, A. Chambouvet, D. S. Milner, V. Attah, R. Terrado, C. Lovejoy, H. Moreau, A. E. Santoro, É. Derelle, T. A. Richards, Host-derived viral transporter protein for nitrogen uptake in infected marine phytoplankton. Proc. Natl. Acad. Sci. 114 (2017).

59. M. T. Cottrell, C. A. Suttle, Dynamics of lytic virus infecting the photosynthetic marine picoflagellate Micromonas pusilla. Limnol. Oceanogr. 40, 730–739 (1995).

60. J. Bull, K. Christensen, C. Scott, B. Jack, C. Crandall, S. Krone, Phage-Bacterial Dynamics with Spatial Structure: Self Organization around Phage Sinks Can Promote Increased Cell Densities. Antibiotics 7, 8 (2018).

61. C. Igler, Phenotypic flux: The role of physiology in explaining the conundrum of bacterial persistence amid phage attack. Virus Evol. 8, veac086 (2022).

62. M. Krasovec, A. Eyre-Walker, N. Grimsley, C. Salmeron, D. Pecqueur, G. Piganeau, S. Sanchez-Ferandin, Fitness Effects of Spontaneous Mutations in Picoeukaryotic Marine Green Algae. G3 GenesGenomesGenetics 6, 2063–2071 (2016).

63. K. P. Burnham, D. R. Anderson, Eds., “Introduction” in Model Selection and Multimodel Inference (Springer New York, New York, NY, 2004; http://link.springer.com/10.1007/978-0-387-22456-5_1), pp. 1–48.

64. K. P. Burnham, D. R. Anderson, Eds., “Information and Likelihood Theory: A Basis for Model Selection and Inference” in Model Selection and Multimodel Inference (Springer New York, New York, NY, 2004; http://link.springer.com/10.1007/978-0-387-22456-5_2), pp. 49–97.

65. S. Yau, G. Caravello, N. Fonvieille, É. Desgranges, H. Moreau, N. Grimsley, Rapidity of Genomic Adaptations to Prasinovirus Infection in a Marine Microalga. Viruses 10, 441 (2018).

66. S. Vacant, L. F. Benites, C. Salmeron, L. Intertaglia, M. Norest, A. Cadoudal, F. Sanchez, C. Caceres, G. Piganeau, Long-Term Stability of Bacterial Associations in a Microcosm of Ostreococcus tauri (Chlorophyta, Mamiellophyceae). Front. Plant Sci. 13, 814386 (2022).

67. S. E. Heath, S. Collins, Mode of resistance to viral lysis affects host growth across multiple environments in the marine picoeukaryote Ostreococcus tauri. Environ. Microbiol. 18, 4628–4639 (2016).

68. S. Heath, K. Knox, P. Vale, S. Collins, Virus Resistance Is Not Costly in a Marine Alga Evolving under Multiple Environmental Stressors. Viruses 9, 39 (2017).

69. L. Listmann, S. Heath, P. F. Vale, C. E. Schaum, S. Collins, Epidemiological dynamics of viral infection in a marine picoeukaryote. Microbiology [Preprint] (2021). 10.1101/2021.03.23.436719.

70. R. Rousseau, C. Cáceres, G. Piganeau, S. Gourbière, How do the molecular determinants of antiviral resistance shape the dynamic of phytoplankton-virus interaction? Ecology [Preprint] (2026). 10.64898/2026.09.29.755341.

71. A. Murray, G. Jackson, Viral dynamics: a model of the effects of size shape, motion and abundance of single-celled olanktonic organisms and other particles. Mar. Ecol. Prog. Ser. 89, 103–116 (1992).

72. B. Chen, Patterns of thermal limits of phytoplankton. J. Plankton Res. 37, 285–292 (2015).

73. J.-B. Guyon, V. Vergé, P. Schatt, J.-C. Lozano, M. Liennard, F.-Y. Bouget, Comparative Analysis of Culture Conditions for the Optimization of Carotenoid Production in Several Strains of the Picoeukaryote Ostreococcus. Mar. Drugs 16, 76 (2018).

74. S. Barton, D. Padfield, A. Masterson, A. Buckling, N. Smirnoff, G. Yvon-Durocher, Comparative experimental evolution reveals species-specific idiosyncrasies in marine phytoplankton adaptation to warming. Glob. Change Biol. 29, 5261–5275 (2023).

75. E. Derelle, S. Yau, H. Moreau, N. H. Grimsley, Prasinovirus Attack of Ostreococcus Is Furtive by Day but Savage by Night. J. Virol. 92, e01703–17 (2018).

76. N. Joffe, C. Kuhlisch, G. Schleyer, N. S. Ahlers, A. Shemi, A. Vardi, Cell-to-cell heterogeneity drives host–virus coexistence in a bloom-forming alga. ISME J. 18, wrae038 (2024).

77. R. Blanc-Mathieu, M. Krasovec, M. Hebrard, S. Yau, E. Desgranges, J. Martin, W. Schackwitz, A. Kuo, G. Salin, C. Donnadieu, Y. Desdevises, S. Sanchez-Ferandin, H. Moreau, E. Rivals, I. V. Grigoriev, N. Grimsley, A. Eyre-Walker, G. Piganeau, Population genomics of picophytoplankton unveils novel chromosome hypervariability. Sci. Adv. 3, e1700239 (2017).

78. C. Bugnot, T. Alioto, F. Cruz Rodriguez, E. Manirakiza, J. Gomez Garrido, M. Gut, S. Yau, G. Piganeau, Interchromosomal translocations and large deletions drive the evolution of the outlier chromosome in the smallest photosynthetic eukaryote. Genome Biol. Evol., evag162 (2026).

79. A. M. Nedelcu, W. W. Driscoll, P. M. Durand, M. D. Herron, A. Rashidi, ON THE PARADIGM OF ALTRUISTIC SUICIDE IN THE UNICELLULAR WORLD. Evolution 65, 3–20 (2011).

80. D. Demory, L. Arsenieff, N. Simon, C. Six, F. Rigaut-Jalabert, D. Marie, P. Ge, E. Bigeard, S. Jacquet, A. Sciandra, O. Bernard, S. Rabouille, A.-C. Baudoux, Temperature is a key factor in Micromonas –virus interactions. ISME J. 11, 601–612 (2017).

81. C. Lønborg, M. Middelboe, C. P. D. Brussaard, Viral lysis of Micromonas pusilla: impacts on dissolved organic matter production and composition. Biogeochemistry 116, 231–240 (2013).

82. C. P. D. Brussaard, G. Bratbak, A.-C. Baudoux, P. Ruardij, Phaeocystis and its interaction with viruses. Biogeochemistry 83, 201–215 (2007).

83. X. Mari, M.-E. Kerros, M. G. Weinbauer, Virus Attachment to Transparent Exopolymeric Particles along Trophic Gradients in the Southwestern Lagoon of New Caledonia. Appl. Environ. Microbiol. 73, 5245–5252 (2007).

84. A. Vardi, L. Haramaty, B. A. S. Van Mooy, H. F. Fredricks, S. A. Kimmance, A. Larsen, K. D. Bidle, Host–virus dynamics and subcellular controls of cell fate in a natural coccolithophore population. Proc. Natl. Acad. Sci. 109, 19327–19332 (2012).

85. C. P. Laber, J. E. Hunter, F. Carvalho, J. R. Collins, E. J. Hunter, B. M. Schieler, E. Boss, K. More, M. Frada, K. Thamatrakoln, C. M. Brown, L. Haramaty, J. Ossolinski, H. Fredricks, J. I. Nissimov, R. Vandzura, U. Sheyn, Y. Lehahn, R. J. Chant, A. M. Martins, M. J. L. Coolen, A. Vardi, G. R. DiTullio, B. A. S. Van Mooy, K. D. Bidle, Coccolithovirus facilitation of carbon export in the North Atlantic. Nat. Microbiol. 3, 537–547 (2018).

86. S. Krishna, V. Peterson, L. Listmann, J. Hinners, Interactive effects of viral lysis and warming in a coastal ocean identified from an idealized ecosystem model. Ecol. Model. 487, 110550 (2024).

87. P. Frémont, S. J. Beckett, D. Demory, E. Carr, C. L. Follett, D. Lindell, D. Talmy, S. Dutkiewicz, J. S. Weitz, Coexistence of Photosynthetic Marine Microorganisms, Viruses and Grazers: Towards Integration in Ocean Ecosystem Models. Environ. Microbiol. 28, e70295 (2026).

