## Supporting Information for "Epidemiological model selection unravels the molecular processes of antiviral resistance from experimental phytoplankton-virus dynamics"

**Supplementary Materials 1. Description of the ACQ, LOSS and structure of the 18 SRIV and 18 SRIVD models of phytoplankton-virus interaction.**

**Table SM1. Description of ACQ and LOSS according to the molecular determinism of the antiviral resistance.** Ext.: extracellular resistance. Int.: complete or partial (with budding) intracellular resistance.

| Determinism | ACQ | ACQ’ | LOSS |
|---|---|---|---|
| Genomic Mutation (GM) | $(b - c\mathrm{N})\mu\mathrm{S}$ | | |
| Random Phenotypic Plasticity (RPP) | $\gamma\mathrm{S}$ | | $\gamma'\mathrm{R}$ |
| Virus-Induced Phenotypic Plasticity (IPP) - Ext. | $\delta_e\beta\mathrm{SV}$ | | $\delta'\mathrm{R}$ |
| Virus-Induced Phenotypic Plasticity (IPP) - Int. | | $\delta_i\mathrm{I}$ | $\delta'\mathrm{R}$ |

System of equations corresponding to the **SRIV models** under the **extracellular** $(\beta' = 0)$ and **intracellular** $(\beta' > 0)$ **resistance mechanisms** and **genomic mutations** (GM) hypotheses without $(\alpha' = 0)$ or with $(\alpha' > 0)$ Programmed Cell Death (PCD);

$$\frac{\mathrm{dS}}{dt} = (b - c\mathrm{N})(1 - \mu)\mathrm{S} - dS - \beta\mathrm{SV} \quad \text{(SM1.1)}$$

$$\frac{\mathrm{dR}}{dt} = (b - c\mathrm{N})\mu\mathrm{S} + \big((1 - x)b - c\mathrm{N}\big)\mathrm{R} - d\mathrm{R} \quad \text{(SM1.2)}$$

$$\frac{\mathrm{dI}}{dt} = \beta\mathrm{SV} - (\alpha + \alpha' + d)\mathrm{I} \quad \text{(SM1.3)}$$

$$\frac{\mathrm{dV}}{dt} = b_v\alpha\mathrm{I} - d_v\mathrm{V} - \beta\mathrm{SV} - \beta'\mathrm{RV} \quad \text{(SM1.4)}$$

where N = S + R + I.

System of equations corresponding to the **SRIV models** under the **mechanism of a partial intracellular resistance associated with budding** and **genomic mutations** (GM) hypotheses without $(\alpha' = 0)$ or with $(\alpha' > 0)$ PCD;

$$\frac{\mathrm{dS}}{\mathrm{d}t} = (b - c\mathrm{N})(1 - \mu)\mathrm{S} - dS - \beta\mathrm{SV} \quad \text{(SM1.5)}$$

$$\frac{\mathrm{dR}}{\mathrm{d}t} = (b - c\mathrm{N})\mu\mathrm{S} + \big((1 - x)b - c\mathrm{N}\big)\mathrm{R} - d\mathrm{R} - \beta'\mathrm{RV} + \psi\mathrm{R^P} \quad \text{(SM1.6)}$$

$$\frac{\mathrm{dR^P}}{\mathrm{d}t} = \beta'\mathrm{RV} + \big((1 - x)b - c\mathrm{N}\big)\mathrm{R^P} - (d + \psi)\mathrm{R^P} \quad \text{(SM1.7)}$$

$$\frac{\mathrm{dI}}{\mathrm{d}t} = \beta\mathrm{SV} - (\alpha + \alpha' + d)\mathrm{I} \quad \text{(SM1.8)}$$

$$\frac{\mathrm{dV}}{\mathrm{d}t} = b_v\alpha\mathrm{I} + b_{vr}\mathrm{R^P} - d_v\mathrm{V} - \beta\mathrm{SV} - \beta'(\mathrm{R} + \mathrm{R^P})\mathrm{V} \quad \text{(SM1.9)}$$

where N = S + R + $\mathrm{R^P}$ + I.

System of equations corresponding to the **SRIVD models** under the **extracellular** ($\beta' = 0$) and **intracellular** ($\beta' > 0$) **resistance mechanisms** and **genomic mutations** (GM) hypotheses without ($\alpha' = 0$) or with ($\alpha' > 0$) PCD;

$$\frac{\mathrm{dS}}{\mathrm{d}t} = (b - c\mathrm{N})(1-\mu)\mathrm{S} - dS - \beta\mathrm{SV} \quad \text{(SM1.10)}$$

$$\frac{\mathrm{dR}}{\mathrm{d}t} = (b - c\mathrm{N})\mu\mathrm{S} + \big((1-x)b - c\mathrm{N}\big)\,\mathrm{R} - d\mathrm{R} \quad \text{(SM1.11)}$$

$$\frac{\mathrm{dI}}{\mathrm{d}t} = \beta\mathrm{SV} - (\alpha + \alpha' + d)\mathrm{I} \quad \text{(SM1.12)}$$

$$\frac{\mathrm{dV}}{\mathrm{d}t} = b_v\alpha\mathrm{I} - d_v\mathrm{V} - \beta\mathrm{SV} - \beta'\mathrm{RV} - \lambda\mathrm{DV} \quad \text{(SM1.13)}$$

$$\frac{\mathrm{dD}}{\mathrm{d}t} = n_{\mathrm{D}}(\alpha + \alpha')\mathrm{I} - \lambda\mathrm{DV} \quad \text{(SM1.14)}$$

where N = S + R + I.

System of equations corresponding to the **SRIVD model** under the **mechanism of a partial intracellular resistance associated with budding** and **genomic mutations** (GM) hypotheses without ($\alpha' = 0$) or with ($\alpha' > 0$) PCD;

$$\frac{\mathrm{dS}}{\mathrm{d}t} = (b - c\mathrm{N})(1-\mu)\mathrm{S} - dS - \beta\mathrm{SV} \quad \text{(SM1.15)}$$

$$\frac{\mathrm{dR}}{\mathrm{d}t} = (b - c\mathrm{N})\mu\mathrm{S} + \big((1-x)b - c\mathrm{N}\big)\mathrm{R} - d\mathrm{R} - \beta'\mathrm{RV} + \psi\mathrm{R^P} \quad \text{(SM1.16)}$$

$$\frac{\mathrm{dR^P}}{\mathrm{d}t} = \beta'\mathrm{RV} + \big((1-x)b - c\mathrm{N}\big)\mathrm{R^P} - (d + \psi)\mathrm{R^P} \quad \text{(SM1.17)}$$

$$\frac{\mathrm{dI}}{\mathrm{d}t} = \beta\mathrm{SV} - (\alpha + \alpha' + d)\mathrm{I} \quad \text{(SM1.18)}$$

$$\frac{\mathrm{dV}}{\mathrm{d}t} = b_v\alpha\mathrm{I} + b_{vr}\mathrm{R^P} - d_v\mathrm{V} - \beta\mathrm{SV} - \beta'(\mathrm{R} + \mathrm{R^P})\mathrm{V} - \lambda\mathrm{DV} \quad \text{(SM1.19)}$$

$$\frac{\mathrm{dD}}{\mathrm{d}t} = n_{\mathrm{D}}(\alpha + \alpha')\mathrm{I} - \lambda\mathrm{DV} \quad \text{(SM1.20)}$$

where N = S + R + $\mathrm{R^P}$ + I.

System of equations corresponding to the **SRIV models** under the **extracellular** ($\beta' = 0$) or **intracellular** ($\beta' > 0$) **resistance mechanisms** and **random phenotypic plasticity** (RPP) hypotheses without ($\alpha' = 0$) or with ($\alpha' > 0$) PCD;

$$\frac{\mathrm{dS}}{\mathrm{d}t} = (b - c\mathrm{N})\mathrm{S} - d\mathrm{S} + \gamma'\mathrm{R} - \gamma\mathrm{S} - \beta\mathrm{SV} \quad \text{(SM1.21)}$$

$$\frac{\mathrm{dR}}{\mathrm{d}t} = \big((1 - x)b - c\mathrm{N}\big)\mathrm{R} - d\mathrm{R} + \gamma\mathrm{S} - \gamma'\mathrm{R} \quad \text{(SM1.22)}$$

$$\frac{\mathrm{dI}}{\mathrm{d}t} = \beta\mathrm{SV} - (\alpha + \alpha' + d)\mathrm{I} \quad \text{(SM1.23)}$$

$$\frac{\mathrm{dV}}{\mathrm{d}t} = b_v\alpha\mathrm{I} - d_v\mathrm{V} - \beta\mathrm{SV} - \beta'\mathrm{RV} \quad \text{(SM1.24)}$$

where N = S + R + I.

System of equations corresponding to the **SRIV model** under the **mechanism of a partial intracellular resistance associated with budding** and **random phenotypic plasticity** (RPP) hypotheses without ($\alpha' = 0$) or with ($\alpha' > 0$) PCD;

$$\frac{\mathrm{dS}}{\mathrm{d}t} = (b - c\mathrm{N})\mathrm{S} - d\mathrm{S} + \gamma'\mathrm{R} - \gamma\mathrm{S} - \beta\mathrm{SV} \quad \text{(SM1.25)}$$

$$\frac{\mathrm{dR}}{\mathrm{d}t} = \big((1 - x)b - c\mathrm{N}\big)\mathrm{R} - d\mathrm{R} + \gamma\mathrm{S} - \gamma'\mathrm{R} - \beta'\mathrm{RV} + \psi\mathrm{R}^{\mathrm{P}} \quad \text{(SM1.26)}$$

$$\frac{\mathrm{dR}^{\mathrm{P}}}{\mathrm{d}t} = \beta'\mathrm{RV} + \big((1 - x)b - c\mathrm{N}\big)\mathrm{R}^{\mathrm{P}} - (d + \psi)\mathrm{R}^{\mathrm{P}} \quad \text{(SM1.27)}$$

$$\frac{\mathrm{dI}}{\mathrm{d}t} = \beta\mathrm{SV} - (\alpha + \alpha' + d)\mathrm{I} \quad \text{(SM1.28)}$$

$$\frac{\mathrm{dV}}{\mathrm{d}t} = b_v\alpha\mathrm{I} + b_{vr}\mathrm{R}^{\mathrm{P}} - d_v\mathrm{V} - \beta\mathrm{SV} - \beta'(\mathrm{R} + \mathrm{R}^{\mathrm{P}})\mathrm{V} \quad \text{(SM1.29)}$$

where N = S + R + $\mathrm{R}^{\mathrm{P}}$ + I.

System of equations corresponding to the **SRIVD models** under the **extracellular** ($\beta' = 0$) or **intracellular** ($\beta' > 0$) **resistance mechanisms** and **random phenotypic plasticity** (RPP) hypotheses without ($\alpha' = 0$) or with ($\alpha' > 0$) PCD;

$$\frac{\mathrm{dS}}{\mathrm{d}t} = (b - c\mathrm{N})\mathrm{S} - d\mathrm{S} + \gamma'\mathrm{R} - \gamma\mathrm{S} - \beta\mathrm{SV} \quad \text{(SM1.30)}$$

$$\frac{\mathrm{dR}}{\mathrm{d}t} = \big((1 - x)b - c\mathrm{N}\big)\mathrm{R} - d\mathrm{R} + \gamma\mathrm{S} - \gamma'\mathrm{R} \quad \text{(SM1.31)}$$

$$\frac{\mathrm{dI}}{\mathrm{d}t} = \beta\mathrm{SV} - (\alpha + \alpha' + d)\mathrm{I} \quad \text{(SM1.32)}$$

$$\frac{\mathrm{dV}}{\mathrm{d}t} = b_v\alpha \mathrm{I} - d_v\mathrm{V} - \beta\mathrm{SV} - \beta'\mathrm{RV} - \lambda\mathrm{DV} \quad \text{(SM1.33)}$$

$$\frac{\mathrm{dD}}{\mathrm{d}t} = n_{\mathrm{D}}(\alpha + \alpha')\mathrm{I} - \lambda\mathrm{DV} \quad \text{(SM1.34)}$$

where N = S + R + I.

System of equations corresponding to the **SRIVD model** under the **mechanism of a partial intracellular resistance associated with budding** and **random phenotypic plasticity** (RPP) hypotheses without ($\alpha' = 0$) or with ($\alpha' > 0$) PCD;

$$\frac{\mathrm{dS}}{\mathrm{d}t} = (b - c\mathrm{N})\mathrm{S} - d\mathrm{S} + \gamma'\mathrm{R} - \gamma\mathrm{S} - \beta\mathrm{SV} \quad \text{(SM1.35)}$$

$$\frac{\mathrm{dR}}{\mathrm{d}t} = \big((1 - x)b - c\mathrm{N}\big)\mathrm{R} - d\mathrm{R} + \gamma\mathrm{S} - \gamma'\mathrm{R} - \beta'\mathrm{RV} + \psi\mathrm{R}^{\mathrm{P}} \quad \text{(SM1.36)}$$

$$\frac{\mathrm{dR}^{\mathrm{P}}}{\mathrm{d}t} = \beta'\mathrm{RV} + \big((1 - x)b - c\mathrm{N}\big)\mathrm{R}^{\mathrm{P}} - (d + \psi)\mathrm{R}^{\mathrm{P}} \quad \text{(SM1.37)}$$

$$\frac{\mathrm{dI}}{\mathrm{d}t} = \beta\mathrm{SV} - (\alpha + \alpha' + d)\mathrm{I} \quad \text{(SM1.38)}$$

$$\frac{\mathrm{dV}}{\mathrm{d}t} = b_v\alpha\mathrm{I} + b_{vr}\mathrm{R}^{\mathrm{P}} - d_v\mathrm{V} - \beta\mathrm{SV} - \beta'(\mathrm{R} + \mathrm{R}^{\mathrm{P}})\mathrm{V} - \lambda\mathrm{DV} \quad \text{(SM1.39)}$$

$$\frac{\mathrm{dD}}{\mathrm{d}t} = n_{\mathrm{D}}(\alpha + \alpha')\mathrm{I} - \lambda\mathrm{DV} \quad \text{(SM1.40)}$$

where N = S + R + $\mathrm{R}^{\mathrm{P}}$ + I.

System of equations corresponding to the **SRIV models** under the **extracellular resistance mechanism** and **virus-induced phenotypic plasticity** (IPP) hypotheses, without ($\alpha' = 0$) or with ($\alpha' > 0$) PCD;

$$\frac{\mathrm{dS}}{\mathrm{d}t} = (b - c\mathrm{N})\mathrm{S} - d\mathrm{S} + \delta'\mathrm{R} - \beta\mathrm{SV} \quad \text{(SM1.41)}$$

$$\frac{\mathrm{dR}}{\mathrm{d}t} = \big((1 - x)b - c\mathrm{N}\big)\mathrm{R} - d\mathrm{R} + \delta_e\beta\mathrm{SV} - \delta'\mathrm{R} \quad \text{(SM1.42)}$$

$$\frac{\mathrm{dI}}{\mathrm{d}t} = (1 - \delta_e)\beta\mathrm{SV} - (\alpha + \alpha' + d)\mathrm{I} \quad \text{(SM1.43)}$$

$$\frac{\mathrm{dV}}{\mathrm{d}t} = b_v\alpha\mathrm{I} - d_v\mathrm{V} - (1 - \delta_e)\beta\mathrm{SV} \quad \text{(SM1.44)}$$

where N = S + R + I.

System of equations corresponding to the **SRIV models** under the **intracellular resistance mechanism** and **virus-induced phenotypic plasticity** (IPP) hypotheses, without ($\alpha' = 0$) or with ($\alpha' > 0$) PCD;

$$\frac{\mathrm{dS}}{\mathrm{d}t} = (b - c\mathrm{N})\mathrm{S} - d\mathrm{S} + \delta'\mathrm{R} - \beta\mathrm{SV} \qquad \text{(SM1.45)}$$

$$\frac{\mathrm{dR}}{\mathrm{d}t} = \big((1-x)b - c\mathrm{N}\big)\mathrm{R} - d\mathrm{R} + \delta_i\mathrm{I} - \delta'\mathrm{R} \qquad \text{(SM1.46)}$$

$$\frac{\mathrm{dI}}{\mathrm{d}t} = \beta\mathrm{SV} - (\alpha + \alpha' + d + \delta_i)\mathrm{I} \qquad \text{(SM1.47)}$$

$$\frac{\mathrm{dV}}{\mathrm{d}t} = b_v\alpha\mathrm{I} - d_v\mathrm{V} - \beta\mathrm{SV} - \beta'\mathrm{RV} \qquad \text{(SM1.48)}$$

where N = S + R + I.

System of equations corresponding to the **SRIV model** under the **mechanism of a partial intracellular resistance associated with budding** and **virus-induced phenotypic plasticity** (IPP) hypotheses without ($\alpha' = 0$) or with ($\alpha' > 0$) PCD;

$$\frac{\mathrm{dS}}{\mathrm{d}t} = (b - c\mathrm{N})\mathrm{S} - d\mathrm{S} + \delta'\mathrm{R} - \beta\mathrm{SV} \qquad \text{(SM1.49)}$$

$$\frac{\mathrm{dR}}{\mathrm{d}t} = \big((1-x)b - c\mathrm{N}\big)\mathrm{R} - d\mathrm{R} - \delta'\mathrm{R} - \beta'\mathrm{RV} + \psi\mathrm{R^P} \qquad \text{(SM1.50)}$$

$$\frac{\mathrm{dR^P}}{\mathrm{d}t} = \delta_i\mathrm{I} + \beta'\mathrm{RV} + \big((1-x)b - c\mathrm{N}\big)\mathrm{R^P} - (d + \psi)\mathrm{R^P} \qquad \text{(SM1.51)}$$

$$\frac{\mathrm{dI}}{\mathrm{d}t} = \beta\mathrm{SV} - (\alpha + \alpha' + d + \delta_i)\mathrm{I} \qquad \text{(SM1.52)}$$

$$\frac{\mathrm{dV}}{\mathrm{d}t} = b_v\alpha\mathrm{I} + b_{vr}\mathrm{R^P} - d_v\mathrm{V} - \beta\mathrm{SV} - \beta'(\mathrm{R} + \mathrm{R^P})\mathrm{V} \qquad \text{(SM1.53)}$$

where N = S + R + $\mathrm{R^P}$ + I.

System of equations corresponding to the **SRIVD models** under the **extracellular resistance mechanism** and **virus-induced phenotypic plasticity** (IPP) hypotheses without ($\alpha' = 0$) or with ($\alpha' > 0$) PCD;

$$\frac{\mathrm{dS}}{\mathrm{d}t} = (b - c\mathrm{N})\mathrm{S} - d\mathrm{S} + \delta'\mathrm{R} - \beta\mathrm{SV} \qquad \text{(SM1.54)}$$

$$\frac{\mathrm{dR}}{\mathrm{d}t} = \big((1-x)b - c\mathrm{N}\big)\mathrm{R} - d\mathrm{R} + \delta_e\beta\mathrm{SV} - \delta'\mathrm{R} \qquad \text{(SM1.55)}$$

$$\frac{\mathrm{dI}}{\mathrm{d}t} = (1 - \delta_e)\beta\mathrm{SV} - (\alpha + \alpha' + d)\mathrm{I} \qquad \text{(SM1.56)}$$

$$\frac{\mathrm{dV}}{\mathrm{d}t} = b_v\alpha\mathrm{I} - d_v\mathrm{V} - (1-\delta_e)\beta\mathrm{SV} - \beta'\mathrm{RV} - \lambda\mathrm{DV} \quad \text{(SM1.57)}$$

$$\frac{\mathrm{dD}}{\mathrm{d}t} = n_\mathrm{D}(\alpha + \alpha')\mathrm{I} - \lambda\mathrm{DV} \quad \text{(SM1.58)}$$

where N = S + R + I.

System of equations corresponding to the **SRIVD models** under the **intracellular resistance mechanism** and **virus-induced phenotypic plasticity** (IPP) hypotheses without $(\alpha' = 0)$ or with $(\alpha' > 0)$ PCD;

$$\frac{\mathrm{dS}}{\mathrm{d}t} = (b - c\mathrm{N})\mathrm{S} - d\mathrm{S} + \delta'\mathrm{R} - \beta\mathrm{SV} \quad \text{(SM1.59)}$$

$$\frac{\mathrm{dR}}{\mathrm{d}t} = \big((1-x)b - c\mathrm{N}\big)\mathrm{R} - d\mathrm{R} + \delta_i\mathrm{I} - \delta'\mathrm{R} \quad \text{(SM1.60)}$$

$$\frac{\mathrm{dI}}{\mathrm{d}t} = \beta\mathrm{SV} - (\alpha + \alpha' + d + \delta_i)\mathrm{I} \quad \text{(SM1.61)}$$

$$\frac{\mathrm{dV}}{\mathrm{d}t} = b_v\alpha\mathrm{I} - d_v\mathrm{V} - \beta\mathrm{SV} - \beta'\mathrm{RV} - \lambda\mathrm{DV} \quad \text{(SM1.62)}$$

$$\frac{\mathrm{dD}}{\mathrm{d}t} = n_\mathrm{D}(\alpha + \alpha')\mathrm{I} - \lambda\mathrm{DV} \quad \text{(SM1.63)}$$

where N = S + R + I.

System of equations corresponding to the **SRIVD model** under the **mechanism of a partial intracellular resistance associated with budding** and **virus-induced phenotypic plasticity** (IPP) without $(\alpha' = 0)$ or with $(\alpha' > 0)$ PCD;

$$\frac{\mathrm{dS}}{\mathrm{d}t} = (b - c\mathrm{N})\mathrm{S} - d\mathrm{S} + \delta'\mathrm{R} - \beta\mathrm{SV} \quad \text{(SM1.64)}$$

$$\frac{\mathrm{dR}}{\mathrm{d}t} = \big((1-x)b - c\mathrm{N}\big)\mathrm{R} - d\mathrm{R} - \delta'\mathrm{R} - \beta'\mathrm{RV} + \psi\mathrm{R^P} \quad \text{(SM1.65)}$$

$$\frac{\mathrm{dR^P}}{\mathrm{d}t} = \delta_i\mathrm{I} + \beta'\mathrm{RV} + \big((1-x)b - c\mathrm{N}\big)\mathrm{R^P} - (d + \psi)\mathrm{R^P} \quad \text{(SM1.66)}$$

$$\frac{\mathrm{dI}}{\mathrm{d}t} = \beta\mathrm{SV} - (\alpha + \alpha' + d + \delta_i)\mathrm{I} \quad \text{(SM1.67)}$$

$$\frac{\mathrm{dV}}{\mathrm{d}t} = b_v\alpha\mathrm{I} + b_{vr}\mathrm{R^P} - d_v\mathrm{V} - \beta\mathrm{SV} - \beta'(\mathrm{R} + \mathrm{R^P})\mathrm{V} - \lambda\mathrm{DV} \quad \text{(SM1.68)}$$

$$\frac{\mathrm{dD}}{\mathrm{d}t} = n_\mathrm{D}(\alpha + \alpha')\mathrm{I} - \lambda\mathrm{DV} \quad \text{(SM1.68)}$$

where N = S + R + $\mathrm{R^P}$ + I.

### Supplementary Materials 2. Estimates of the aggregation parameters $r_P$ and $r_V$ of the negative binomial distributions accounting for the overdispersion in phytoplankton and virus abundance data.

We estimated the value for the aggregation parameters $r_P$ and $r_V$ prior to models fitting while considering that the over-dispersed distribution accounting for the variability of the data is independent of the model being fitted. This allowed to reduce the dimension of the parameter space to be explored while fitting each of the 36 proposed models. To do so, we first reproduced the average dynamics of the phytoplankton and virus kinetics through polynomial regressions (Figure SM1). We then found the value of the aggregation parameters best reproducing dispersal around the average dynamic by maximizing the following log-likelihood function for *O. tauri* (SM2.1) and OtV5 (SM2.2);

$$LLH(O_P|r_\mathrm{P}) = \sum_{i=4}^{n}\left[\sum_{j=1}^{m} \log \mathrm{p}\big(\mathrm{P}(t_j) = \mathrm{O}_P(t_j)\big|r_P\big)\right] \qquad \text{(SM2.1)}$$

where, for the data subset $i$, $\mathrm{P}(t_j)$ is the statistical variable corresponding to the abundance of phytoplankton at time $t_j$ (in hours) in the polynomial regression and $\mathrm{O}_P(t_j)$ stands for the observed abundance at that time of the experiment. Probabilities $\mathrm{p}\big(\mathrm{P}(t_j) = \mathrm{O}_P(t_j)\big|r_\mathrm{P}\big)$ were calculated according to the negative binomial distribution accounting for the overdispersion in phytoplankton abundance (*62*) and according to the aggregation parameter $r_P$ being tested.

The same approach was then independently intended for the virus dynamics, by maximizing the following log-likelihood function (SM2.2);

$$LLH(O_V|r_\mathrm{V}) = \sum_{i=3}^{n}\left[\sum_{j=1}^{m} \log \mathrm{p}\big(\mathrm{V}(t_j) = \mathrm{O}_V(t_j)\big|r_V\big)\right] \qquad \text{(SM2.2)}$$

This approach provided us with estimates of $r_P$ equals to 94.6 and of $r_V$ equals to 36.1 that fitted well the distribution of phytoplankton and virus data (Figure SM1).

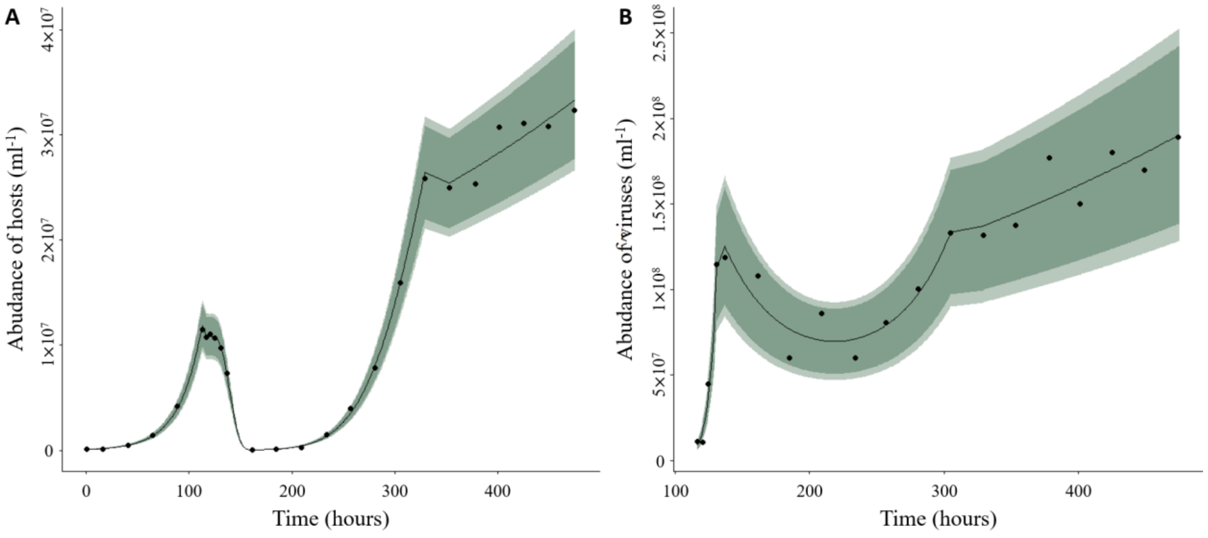


**Figure SM1. Experimental dynamic of *O. tauri*–OtV5 interaction and polynomial predictions of their average dynamics.** The observed variations in (A) *O. tauri* and (B) OtV5 abundances used for polynomial regression with negative binomial distributions. The black curves show the prediction of the average dynamics of the phytoplankton and virus populations. The light and dark green areas correspond to the 95% and 99% confidence intervals derived from the Negative Binomial distributions with aggregation parameter estimates of $r_P$ (94.6) and $r_V$ (36.1).

### Supplementary Materials 3. Complete ranking of the 36 models in the multiple hypotheses testing through model selection.

**Table SM1. Summary of the model selection analysis of the 36 competing models.** The column 'Model' gives the ranking, the type (SRIV/SRIVD), the resistance hypotheses (about determinism and mechanism), and the number of parameters for a given model *i*. There, RP stands for (partial intracellular) Resistant Producers (through budding). The outcomes of each model fitting to the experimental kinetics are provided in the other columns; the maximum likelihood $(L_i)$, the corrected AIC $(AIC_{c_i})$, the difference between the model's AICc and the AICc of the best model $(\Delta_i(AIC_c))$, the weights of Akaike $(w_i(AIC_c))$, and the values of $R_h^2$ and $R_v^2$ that stand for the coefficients of determination derived from the linear regression between observed and predicted abundances. The values of $R_h^2$ and $R_v^2$ appearing in black and grey correspond to model fitting and validation, respectively.

| Model | | | | $n$ | $L_i$ | $AIC_{ci}$ | $\Delta_i(AIC_c)$ | $w_i(AIC_c)$ | $R_h^2$ | $R_v^2$ |
|---|---|---|---|---|---|---|---|---|---|---|
| 1 | SRIVD | IPP | RP | 15 | -599.0113 | 1252 | 0 | 0.65 | 0.98/0.99 | 0.94/0.90 |
| 2 | SRIVD | RPP | RP | 15 | -599.7769 | 1253.6 | 1.6 | 0.29 | 0.98/0.99 | 0.94/0.90 |
| 3 | SRIVD | IPP | RP PCD | 16 | -599.0113 | 1258.7 | 6.7 | 0.02 | 0.98/0.99 | 0.94/0.90 |
| 4 | SRIVD | RPP | RP PCD | 16 | -599.7625 | 1260.2 | 8.2 | 0.01 | 0.98/0.99 | 0.94/0.90 |
| 5 | SRIV | IPP | RP | 13 | -609.0269 | 1260.6 | 8.6 | 0.01 | 0.98/0.99 | 0.83/0.79 |
| 6 | SRIV | RPP | RP | 13 | -609.6034 | 1261.8 | 9.8 | 0.01 | 0.98/0.99 | 0.83/0.79 |
| 7 | SRIVD | GM | RP | 14 | -607.1215 | 1262.2 | 10.2 | < 0.01 | 0.98/0.99 | 0.85/0.85 |
| 8 | SRIVD | RPP | Extracellular | 12 | -612.7540 | 1263.1 | 11.1 | < 0.01 | 0.98/0.99 | 0.85/0.83 |
| 9 | SRIVD | IPP | Extracellular | 12 | -612.7643 | 1263.1 | 11.1 | < 0.01 | 0.98/0.99 | 0.85/0.82 |
| 10 | SRIV | IPP | RP PCD | 14 | -608.8562 | 1265.7 | 13.7 | < 0.01 | 0.98/0.99 | 0.83/0.79 |
| 11 | SRIV | RPP | RP PCD | 14 | -609.4212 | 1266.8 | 14.8 | 0 | 0.98/0.99 | 0.84/0.79 |
| 12 | SRIVD | RPP | Extracellular PCD | 13 | -612.7540 | 1268.1 | 16.1 | 0 | 0.98/0.99 | 0.85/0.83 |
| 13 | SRIVD | RPP | Intracellular | 13 | -612.7540 | 1268.1 | 16.1 | 0 | 0.98/0.99 | 0.85/0.82 |
| 14 | SRIVD | IPP | Extracellular PCD | 13 | -612.7613 | 1268.1 | 16.1 | 0 | 0.98/0.99 | 0.84/0.82 |
| 15 | SRIVD | GM | RP PCD | 15 | -607.0828 | 1268.2 | 16.2 | 0 | 0.98/0.99 | 0.84/0.85 |
| 16 | SRIVD | IPP | Intracellular | 13 | -612.8724 | 1268.3 | 16.3 | 0 | 0.98/0.99 | 0.83/0.81 |
| 17 | SRIVD | RPP | Intracellular PCD | 14 | -612.7540 | 1273.5 | 21.5 | 0 | 0.98/0.99 | 0.85/0.83 |
| 18 | SRIVD | IPP | Intracellular PCD | 14 | -612.8724 | 1273.7 | 21.7 | 0 | 0.98/0.99 | 0.83/0.81 |
| 19 | SRIVD | GM | Extracellular | 11 | -630.3602 | 1293.7 | 41.7 | 0 | 0.98/0.99 | 0.48/0.43 |
| 20 | SRIVD | GM | Extracellular PCD | 12 | -629.1352 | 1295.8 | 43.8 | 0 | 0.98/0.99 | 0.48/0.43 |
| 21 | SRIVD | GM | Intracellular | 12 | -630.3602 | 1298.3 | 46.3 | 0 | 0.98/0.99 | 0.48/0.43 |
| 22 | SRIVD | GM | Intracellular PCD | 13 | -629.1352 | 1300.8 | 48.8 | 0 | 0.98/0.99 | 0.48/0.43 |
| 23 | SRIV | GM | RP | 12 | -633.4420 | 1304.4 | 52.4 | 0 | 0.97/0.99 | 0.58/0.58 |
| 24 | SRIV | GM | RP PCD | 13 | -632.1289 | 1306.8 | 54.8 | 0 | 0.98/0.99 | 0.66/0.65 |
| 25 | SRIV | RPP | Extracellular | 10 | -643.4716 | 1315.7 | 63.7 | 0 | 0.98/0.99 | 0.58/0.57 |
| 26 | SRIV | IPP | Extracellular | 10 | -643.4768 | 1315.8 | 63.8 | 0 | 0.98/0.99 | 0.58/0.57 |
| 27 | SRIV | IPP | Extracellular PCD | 11 | -642.2210 | 1317.4 | 65.4 | 0 | 0.98/0.99 | 0.60/0.59 |
| 28 | SRIV | RPP | Extracellular PCD | 11 | -642.2341 | 1317.5 | 65.5 | 0 | 0.98/0.99 | 0.60/0.58 |
| 29 | SRIV | RPP | Intracellular | 11 | -643.4716 | 1319.9 | 67.9 | 0 | 0.98/0.99 | 0.58/0.57 |
| 30 | SRIV | IPP | Intracellular | 11 | -643.9898 | 1321 | 69 | 0 | 0.98/0.99 | 0.57/0.56 |
| 31 | SRIV | RPP | Intracellular PCD | 12 | -642.2341 | 1322 | 70 | 0 | 0.98/0.99 | 0.60/0.58 |
| 32 | SRIV | IPP | Intracellular PCD | 12 | -643.4725 | 1324.5 | 72.5 | 0 | 0.98/0.99 | 0.59/0.58 |
| 33 | SRIV | GM | Extracellular | 9 | -652.9136 | 1330.8 | 78.8 | 0 | 0.98/0.99 | 0.41/0.37 |
| 34 | SRIV | GM | Extracellular PCD | 10 | -651.6863 | 1332.2 | 80.2 | 0 | 0.98/0.99 | 0.41/0.37 |
| 35 | SRIV | GM | Intracellular | 10 | -652.9136 | 1334.6 | 82.6 | 0 | 0.98/0.99 | 0.41/0.37 |
| 36 | SRIV | GM | Intracellular PCD | 11 | -651.6863 | 1336.4 | 84.4 | 0 | 0.98/0.99 | 0.41/0.37 |
